# Multi-timescale dynamics in descending pain modulation

**DOI:** 10.64898/2025.12.15.693905

**Authors:** Carl Ashworth, Melissa Martenson, Zhigang Shi, Caitlynn C. De Preter, Mary Heinricher, Flavia Mancini

## Abstract

Descending pain-modulatory circuits must coordinate rapid nociceptive responses with slower changes in physiological state, yet how their activity is organized across time remains poorly understood. We investigated this question in physiologically identified pain-modulating neurons of the rostral ventromedial medulla (RVM). The RVM contains ON-and OFF-cells that exert descending control over spinal nociceptive transmission, regulating pain sensitivity and behaviors. By integrating neuronal recordings with probabilistic modeling, we show that unstimulated and stimulus-driven conditions give rise to distinct timescales of ON-and OFF-cell dynamics. During noxious stimulation, we find that population responses undergo rapid activation (*<* 1 s) followed by superimposed slow (∼ 100 s) and fast (*<* 10 s) recovery dynamics. In the absence of stimulation, the same neurons exhibit slow quasi-periodic fluctuations in firing activity with a characteristic timescale of approximately five minutes. Gaussian-process models show that these slow dynamics are statistically predictable from past activity, indicating structured temporal organization beyond stimulus-evoked responses. Taken together, these results indicate that descending pain-control circuits exhibit structured dynamics encompassing rapid pain-related signaling and slower fluctuations consistent with ongoing physiological state.

**Significance:** Pain regulation requires coordination between rapid defensive responses and slower changes in physiological state, yet how these processes are integrated within descending pain-modulatory circuits remains unclear. We show that neurons in a key brainstem pain-control center, the rostral ventromedial medulla, operate across multiple timescales. Using neuronal recordings and computational modeling, we find that these neurons exhibit both fast responses to painful stimuli and slow, structured fluctuations in ongoing activity. These results demonstrate that descending pain-modulatory neurons exhibit temporal organization beyond immediate reflex-linked responses and provide a quantitative framework for investigating how descending pain modulation is regulated over time.

## Introduction

Neural circuits that regulate behavior must integrate rapid sensorimotor responses with slower fluctuations in internal physiological state. Reflexive reactions unfold within milliseconds, whereas sensory gain and behavioral sensitivity can vary over seconds to minutes depending on arousal, context, and physiological state. In the nociceptive system, this coordination across timescales is particularly important because descending brainstem pathways dynamically regulate pain sensitivity and are major targets of pharmacological and neuromodulatory interventions. However, despite the central role of these circuits in controlling pain and mediating therapeutic effects, how neural activity within descending pain-control pathways is organized across multiple timescales remains poorly understood. Understanding these temporal dynamics is therefore important for interpreting how pharmacological and neuromodulatory interventions engage descending pain-control circuits and may ultimately help inform the development and optimization of such therapies.

A central component of this descending pain-control system is the rostral ventromedial medulla (RVM), a key node in a modulatory pathway that regulates spinal nociceptive processing ^1,2^. Through direct projections to the dorsal horn, the RVM exerts potent inhibitory and facilitatory control over nociceptive transmission ^3–9^, dynamically adjusting pain sensitivity according to behavioral and physiological context. This descending modulation is mediated by two principal classes of RVM neurons: ON-and OFF-cells. Increased ON-cell activity facilitates nociceptive transmission and promotes hyperalgesia, whereas activation of OFF-cells suppresses nociceptive signaling and produces hypoalgesia. During noxious stimulation, ON-cells increase firing immediately prior to withdrawal responses, while OFF-cells pause, providing a physiological signature for identifying both populations ^10^ (Figure 1). Cells that do not exhibit stimulus-linked modulation are typically classified as NEUTRAL-cells. Together, these populations contribute to a descending control system that dynamically regulates nociceptive gain. Experimental activation of OFF-and ON-cells respectively results in hypoalgesia and hyperalgesia ^6–9,11–14^. Furthermore, inactivation of ON-cells attenuates hypersensitivity in persistent pain models ^15–20^. More generally, periods of ON-cell dominance have been shown to be associated with relative hypersensitivity compared to periods of OFF-cell dominance ^21^. Therefore, RVM function cannot be understood solely through reflex-linked ON-and OFF-cell responses; ongoing fluctuations in these populations may also contribute to descending pain modulation.

**Figure 1:**
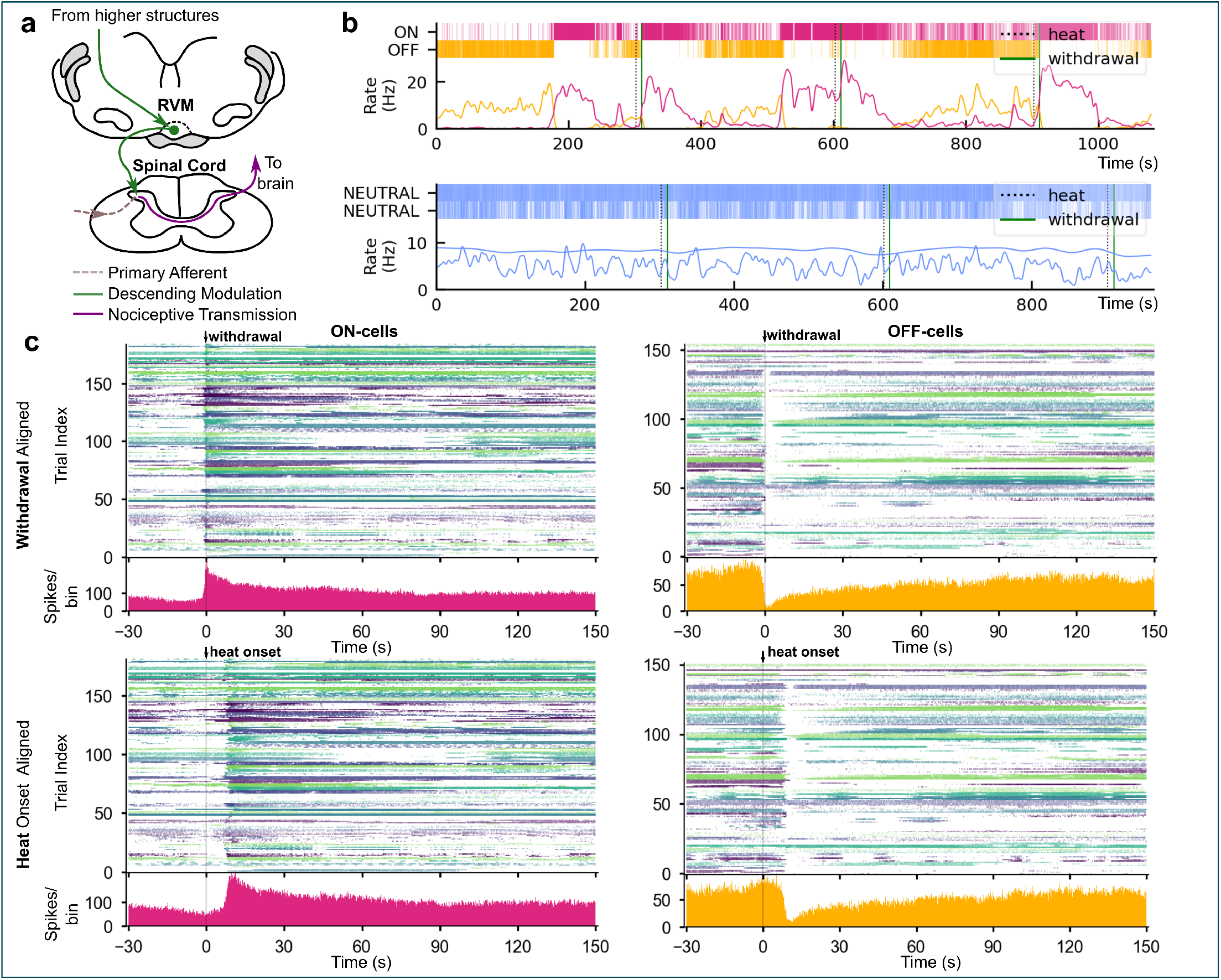
Stimulation-evoked activity of ON-, OFF-, and NEUTRAL-cells in the RVM. **a**, Schematic of the RVM and its descending projections to the spinal dorsal horn, illustrating the modulatory roles of ON-and OFF-cells in nociceptive processing. **b**, Representative spike rasters (top) and rate plots (bottom) from simultaneously recorded ON-, OFF-, and NEUTRAL-cells during repeated noxious heat trials. ON-and OFF-cells exhibit the canonical reciprocal dynamics around paw withdrawal (solid green line), whereas NEUTRAL-cells show no clear modulation. Notably, alternating dominance of ON-and OFF-cells is also observed during inter-trial intervals, demonstrating ongoing activity that is not directly linked to noxious stimulation. **c**, Population peri-stimulus time histograms (PSTHs) for ON-and OFF-cells, aligned either to the time of paw withdrawal (top) or to the heat stimulus onset (bottom). Population firing is sharply time-locked to the withdrawal reflex but not to the onset of heat application. Withdrawal-aligned PSTHs show steeper slopes and no delay compared to heat-aligned PSTHs, consistent with the nocifensive response, rather than the stimulus as such, being the primary driver of RVM activity.

Consistent with this possibility, several features of RVM organization suggest that its function extends beyond reflex-linked modulation. The RVM integrates bottom-up sensory input from the spinal cord with sustained top-down influences from the periaqueductal gray, hypothalamus, and other homeostatic centers ^22–24^. Studies comparing direct and indirect recruitment of spinal and trigeminal inputs through the parabrachial complex further show that RVM neurons can exhibit both transient and prolonged responses depending on the route of activation, suggesting that they integrate nociceptive signals across distinct temporal scales ^25^. Early electrophysiological recordings demonstrated that ON-and OFF-cells show broadly anticorrelated fluctuations even in the absence of sensory stimulation ^26^, and that these fluctuations are associated with measurable changes in nociceptive threshold ^21^. However, these observations were largely qualitative, and neither the temporal structure nor the predictability of evoked and ongoing RVM activity has been quantitatively characterized. In particular, it remains unknown whether ongoing fluctuations exhibit reproducible temporal organization or simply reflect stochastic variability and changing physiological state.

We hypothesized that RVM neurons operate across multiple temporal scales, integrating fast responses associated with reflex-linked control with slower fluctuations consistent with ongoing network or state-dependent modulation. To test this idea, we combined neuronal recordings from identified ON-, OFF-, and NEUTRAL-cells with probabilistic modeling to characterize both stimulus-evoked and ongoing activity in the RVM using a robust lightly anesthetized rodent model in which activation of OFF-cells has been demonstrated to produce hypoalgesia and activation of ON-cells to produce hyperalgesia ^11,12,15,27^. This preparation is particularly suited to stable single-unit recordings from the RVM because the region is deep and physiologically identified ON-and OFF-cells are relatively sparse, making prolonged maintenance of contact with individual neurons difficult in awake animals. Light anesthesia suppresses spontaneous movement while preserving nocifensive withdrawal responses and the canonical physiological signatures used to identify ON-, OFF-, and NEUTRAL-cells ^10,28^. Although anesthesia may influence slow network dynamics, this preparation therefore provides the recording stability required to characterize the activity of identified RVM neurons over extended periods.

Our analyses reveal multi-timescale population dynamics in which fast, multi-phase responses to noxious stimuli unfold over seconds to tens of seconds, while slower quasi-periodic fluctuations in ongoing activity occur over minutes. These slow quasi-periodic fluctuations, observed preferentially in ON-and OFF-cells, were statistically predictable from past activity. The present data do not establish whether these dynamics originate within the RVM or reflect interactions with broader physiological or network processes. Together, these findings show that descending pain-modulatory neurons exhibit structured dynamics encompassing multiple temporal scales and provide a quantitative framework for investigating how descending pain modulation is organized across time.

## Results

### Canonical ON/OFF responses to noxious stimulation

To understand how descending pain-modulatory neurons operate across multiple timescales, we first examined the activity of physiologically identified RVM neurons during noxious stimulation, consistent with previous work^10,26,29–31^. We used a well-established lightly anesthetized preparation with extracellular single-unit recordings (see Methods), which preserves reliable nocifensive withdrawal responses while minimizing spontaneous movement, thereby allowing stable long-duration recordings from identified neurons in the deep RVM. Recordings from ON-and OFF-cells revealed structured population responses that extended well beyond the brief epoch surrounding withdrawal behavior (Figure 1). These neurons exhibited multi-phase activity patterns comprising an initial rapid stimulus-linked change in firing followed by prolonged recovery dynamics lasting tens of seconds.

Figure 1b shows representative recordings from an ON-cell and OFF-cell recorded simultaneously in one animal, alongside two NEUTRAL-cells recorded in a second animal. During noxious heat stimulation (vertical dashed line), the ON-cell increased firing in close temporal proximity to paw with-drawal, whereas the OFF-cell exhibited a sharp pause in firing, consistent with canonical reciprocal dynamics. In contrast, NEUTRAL-cells showed no stimulus-associated changes in firing (Supplementary Figure S3), consistent with their canonical lack of nociceptive responsiveness. Notably, spontaneous switches between OFF-and ON-dominant activity were also observed during inter-trial intervals (e.g., at 180 s and 520 s), indicating that the activity of these cell classes is not solely stimulus-evoked.

Population responses are summarized in Figure 1c, which shows peri-stimulus time histograms (PSTHs) for 70 ON-cells and 60 OFF-cells, aligned either to paw withdrawal (Figure 1c, upper) or heat onset (Figure 1c, lower). In total, 185 ON-cell and 155 OFF-cell trials contributed to the withdrawal-aligned PSTHs, while 182 ON-cell and 151 OFF-cell trials contributed to the heat-onset-aligned PSTHs. These PSTHs were computed over a 150-s window and therefore include slightly fewer trials than the Bayesian analyses presented below, which used a shorter analysis window. A detailed breakdown of trials and animals is provided in the Supplementary Information. Population PSTHs were obtained by pooling responses across neurons and animals to estimate the shared temporal dynamics of each physiologically defined cell class. Characterizing neuron-to-neuron variability or population synchrony would require substantially larger numbers of simultaneously recorded neurons than are available in the present dataset.

PSTHs showed a clear lag and shallower slopes when aligned to heat onset compared to withdrawal, confirming that our dataset captures the canonical, reciprocal ON/OFF responses associated with nocifensive reflexes. Notably, ON-and OFF-cell activity also evolved on longer timescales that extended beyond the time of withdrawal, suggesting that the evoked response could not be adequately described by a single recovery timescale. We therefore modeled ON-and OFF-cell population dynamics using Bayesian regression to characterize these response and recovery timescales.

### Evoked ON/OFF activity unfolds over multiple timescales

To characterize the evoked response and recovery dynamics of ON-and OFF-cell populations, we applied a piecewise non-linear Bayesian regression model with a Poisson likelihood to the summed spike counts per time bin across the entire ON-and OFF-cell populations, aligned initially by withdrawal time (Figure 2a; see Methods). We chose a window from 10 seconds before, to 100 seconds post withdrawal (or heat onset, discussed later). This resulted in 195 ON-cell and 170 OFF-cell trials aligned to withdrawal, and 197 ON-cell and 172 OFF-cell trials aligned to heat onset. Cells were not distinguished by their firing rate pre-trial and all trials with a withdrawal were included. We focused on population-level structure rather than animal-or cell-specific effects, given the small number of trials per animal and the absence of a principled basis for differential weighting. Under the assumption of exchangeability, pooling across cells and animals reduces uncorrelated noise and preserves shared population dynamics. We used Bayesian highest density intervals (HDIs), defined as the minimum width Bayesian credible intervals ^32^. A 97% HDI means there is a 97% probability that the unobserved parameter lies within the given interval, given the model and data.

**Figure 2:**
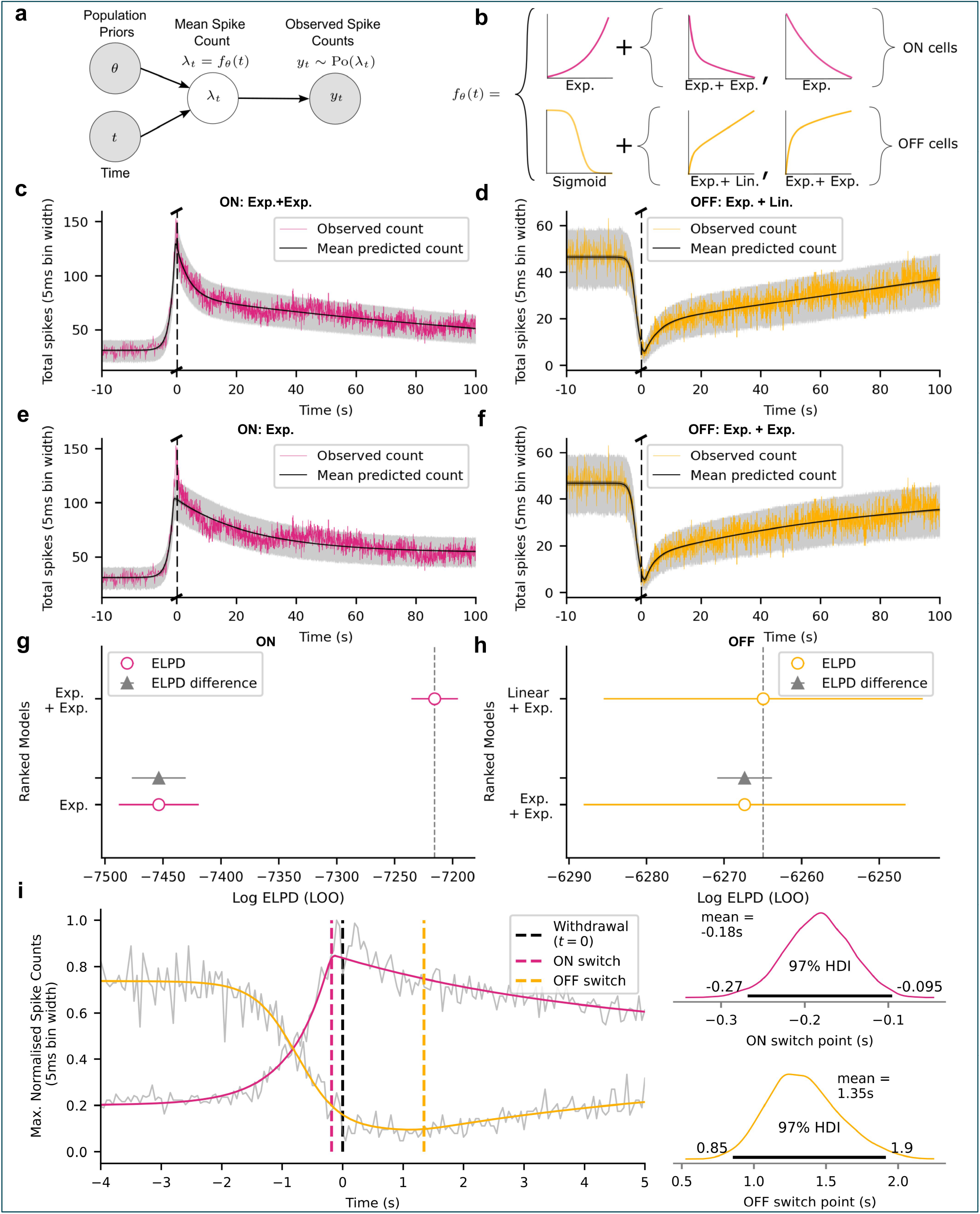
Bayesian modeling of ON-and OFF-cell population dynamics during noxious stimulation. **a**, The probabilistic model used to describe withdrawal-aligned population firing rates. **b**, Candidate response–recovery functions per cell type. For ON-cells, single vs. double exponential (exp.) recovery models were compared; for OFF-cells, double exp. vs. linear + exp. recovery models were evaluated. **c**-**f**, Models were fitted to the population firing rates per cell type and model variant. The single exp. recovery model missed the sharp ON-cell peak (**e**) and the double exp. model underestimated the OFF-cell recovery rate (**f**). **g**-**h**, Expected Log Predictive Density (ELPD) comparison of models, confirming that the double exp. recovery best fit ON-cells, while the linear + exp. model best fit OFF-cells. Whilst the linear + exp. model performed best for OFF-cells, the improvement over the double exp. model was minimal. **i**, Combined ON-and OFF-cell model fits around the withdrawal and population data in gray (left). Posterior switching timepoint estimates (*t*_*s*_) per cell type (right), showing that ON-cells peak slightly before withdrawal, while OFF-cells exhibit post-withdrawal suppression.

For all analyses of evoked activity, we decomposed each trial into two segments: a response phase, defined as the monotonic departure of firing rate from the pre-withdrawal baseline (increase for ON-cells, decrease for OFF-cells), and a recovery phase, defined as the monotonic return toward a post-withdrawal baseline. These were separated by a single transition timepoint *t*_*s*_, corresponding to the extremum (peak for ON-cells, nadir for OFF-cells) before recovery began. A null model assuming a constant mean spike count performed significantly worse than all piecewise models, confirming the presence of both an evoked response and a temporally distinct recovery phase (see Supplementary Figure S8). Since all trials were aligned to withdrawal, variability in the withdrawal latency (time between heat onset and withdrawal) cannot shift post-withdrawal activity into the pre-withdrawal window, making an averaging artifact an unlikely source of the observed recovery dynamics.

The fitted switching timepoint between response and recovery differed for ON-and OFF-cells (Figure 2g): 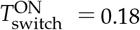 before withdrawal (97% HDI: 0.27–0.095 s before), and 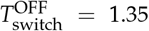 after withdrawal (97% HDI: 0.85–1.9 s after). Thus ON-cells peak slightly before withdrawal, while OFF-cells enter a brief post-withdrawal depression before beginning their recovery phase. This relative timing is visible in previous literature but has not been explicitly modeled ^30^. The response phase was modeled as an exponential rise for ON-cells and a sigmoidal decrease for OFF-cells. We also considered a Gaussian cumulative density function (*er f*) decrease for OFF-cells, which has lighter tails than the sigmoid, and a different biological interpretation (see Methods, Discussion); this fitted significantly worse than the sigmoid across all recovery models (see Supplemental Gaussian CDF Fits, Tables S9 and S10). The recovery phase was parameterized using alternative functional forms: single vs. double exponential recovery for ON-cells, and double exponential vs. linear + exponential recovery for OFF-cells (Figure 2b).

As shown in Figure 2c,g, ON-cell recovery was best captured by a double exponential model, consisting of a superposition of two distinct decay rates (timescales). ON-cell firing rose on a sub-second timescale before paw withdrawal, with time constant *τ*_pre_ = 0.653 s (97% HDI: 0.570–0.739 s). After paw withdrawal, the recovery exhibited both a fast component *τ*_fast_ = 4.157 s (97% HDI: 3.582–4.722 s) and a slow component *τ*_slow_ = 164.223 s (97% HDI: 129.571–199.809 s). A single exponential model (Figure 2e) could not simultaneously capture the sharp evoked peak and the prolonged decay, instead over-and undershooting different portions of the trajectory.

For OFF-cells, recovery was best described by the combined linear and exponential model (Figure 2d,h). OFF-cell firing declined sharply before withdrawal, with a sigmoidal response slope of *k* = 2.852 s^−1^ (97% HDI: 2.333–3.392 s^−1^). The post-withdrawal recovery contained a fast exponential component *τ* = 4.760 s (97% HDI: 3.711–5.861 s), together with a slower linear drift of *k*_linear_= 0.186 counts per second (97% HDI: 0.174–0.196 counts per second). Although the double exponential alternative fit the early part of the recovery (Figure 2f), it failed to asymptote to baseline over the recorded interval, making the linear+exponential model the better descriptor of the OFF-cell dynamics we observed.

We next quantified the timing of the response and recovery phases of the evoked changes in firing rate in terms of percentage deviations from baseline (see Supplemental Figure S1 for full posteriors). OFF-cell population firing had already decreased by 10% from baseline at 1.5 s before the paw withdrawal (97% HDI: 1.746–1.306 s), and ON-cell firing increased by 10% on a similar timescale, at 1.7 s before withdrawal (97% HDI: 1.9–1.5 s), indicating that both populations depart from baseline well before the behavioral response. The two classes also crossed 50% of their total change within a narrow pre-withdrawal window (OFF: 726 ms before; 97% HDI: 810–645 ms; ON: 630 ms before; 97% HDI: 690–570 ms), showing that the dominant portion of the evoked response is temporally locked to the upcoming reflex. Notably, ON-cell firing reached 90% of its peak well before paw withdrawal (240 ms before, 97% HDI: 320–180 ms), while OFF-cell suppression did not reach 90% until shortly after the behavioral response (41 ms post, 97% HDI: 95 ms pre –180 ms post), reflecting a temporal asymmetry in how the two populations complete their evoked transitions.

Comparison of these values to the transition time for OFF-cells described above shows that OFF-cell firing continues to decrease for several seconds post-withdrawal before beginning to recover. At the population level, ON-cell activation both precedes withdrawal and completes its evoked component rapidly, whereas OFF-cell suppression is slower to reach its minimum and persists into the post-withdrawal period. Consistent with this ordering, ON-cell activation reaches its peak before OFF-cell firing reaches full suppression, indicating that, at the population level, peak ON-cell activity precedes maximal OFF-cell suppression. Because this timing was observed in a pseudo-population rather than within simultaneously recorded ON/OFF pairs, it should be interpreted as a population-level temporal organization rather than a claim about pairwise interactions.

We used separate baselines for the pre-and post-withdrawal activity, which allowed us to quantify recovery on the same percentage scale, accounting for any slow modulation of tonic firing relative to the trial length. ON-cells returned to 50% of their post-withdrawal baseline by 15.39 s after the withdrawal (97% HDI: 9.373-22.43 s), but required more than an order of magnitude longer to approach full recovery, reaching 90% of their baseline by 222 s (97% HDI: 183.6-248.8 s). OFF-cells increased back to 50% of their baseline by 18 s post-withdrawal (97% HDI: 15-21 s) but to 90% of their baseline by 83 s (97% HDI: 82-84 s). These timescales are considerably longer than the typical duration of EMG activity ^30^, indicating that the prolonged temporal structure is substantially longer than the duration of the withdrawal reflex itself.

An important consequence of modeling ON-and OFF-cell activity parametrically is that it allows the sharpness of the evoked transitions to be quantified directly, using the slope or time constant of the transitions, rather than inferred qualitatively from peri-stimulus averages (as in Figure 1c) ^33^. For OFF-cells, the slope of the sigmoidal fits to firing-rate decreases was significantly steeper for the withdrawal-aligned data (2.852 total spikes per bin, 97% HDI: 2.333–3.392 spikes per bin), compared to the heat-aligned data (1.747 total spikes per bin, 97% HDI: 1.476–2.041 spikes per bin) (Supplementary Tables S6 and S7). Similarly, ON-cells exhibited faster increases in firing when aligned to withdrawal, with exponential time constants of 0.653 s (97% HDI: 0.570–0.739 s), compared to 1.278 s (97% HDI: 1.129–1.426 s) when aligned to heat onset (Supplementary Tables S3 and S5). In contrast, NEUTRAL-cells (*n* = 32; 69 withdrawal-aligned and 70 heat-aligned trials) exhibited no discernible modulation associated with either heat onset or withdrawal behavior (Supplementary Figure S3). Importantly, these neurons were recorded under the same anesthetic and physiological conditions as the ON and OFF populations, supporting the conclusion that stimulus-linked modulation differed between physiologically identified cell classes.

Taken together, these results show that ON-and OFF-cell dynamics unfold over multiple, cell-type-specific timescales extending well beyond paw withdrawal. This led us to examine whether ON-and OFF-cells exhibit ongoing rhythmic activity in the absence of stimulation, and how such slow oscillations might interact with stimulus-evoked responses.

### RVM ON-and OFF-cells exhibit slow quasiperiodic structure during ongoing activity

To investigate the ongoing activity of RVM neurons, we analyzed ON-, OFF-, and NEUTRAL-cells (*N*_ON_ = 25, *N*_OFF_ = 25, *N*_NEUTRAL_ = 32) recorded in the absence of external stimulation (Figure 3a).

**Figure 3:**
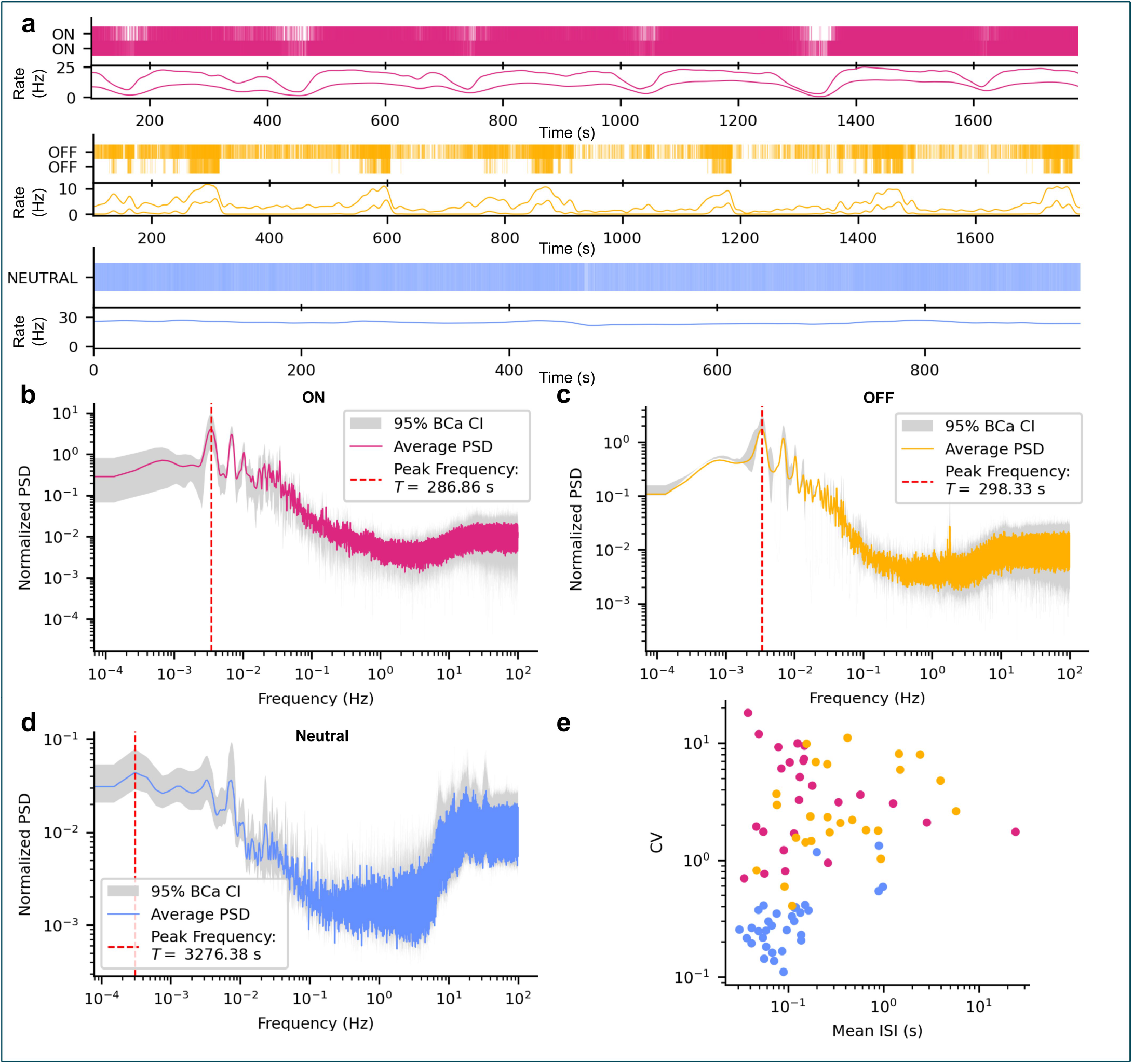
Ongoing activity of RVM neurons reveals distinct low-frequency signatures in ON-and OFF-cells. **a**, Example spike rasters (top) and rate plots (bottom) from simultaneously recorded ON-, OFF-, and NEUTRAL-cells during unstimulated periods. ON-and OFF-cells exhibit irregular, burst-like activity with slow co-varying fluctuations, whereas NEUTRAL-cells fire regularly at a high rate. **b**-**d**, Population-averaged power spectra of ongoing activity for ON-, OFF-, and NEUTRAL-cells. ON-and OFF-cells show prominent low-frequency peaks just below *T* = 300 s (5 min), frequently accompanied by harmonics, while NEUTRAL-cells lack such structure and instead show strong high-frequency components (10–100 Hz). **e**, Coefficient of variation (CV) versus mean interspike interval (ISI) for all recorded cells. NEUTRAL-cells form a distinct cluster characterized by low CV and short ISIs (regular high-rate firing), whereas ON-and OFF-cells show broad CV distributions and longer ISIs, consistent with irregular burst dynamics.

Even in the absence of sensory stimulation, ON-and OFF-cells exhibited slow fluctuations in firing rate (examples in Figure 3a). To determine whether these fluctuations contained reproducible temporal structure, we computed power spectral density (PSD) estimates for each cell class (Figure 3b,c,d). Both ON-and OFF-cells exhibited clear low-frequency spectral peaks, often accompanied by harmonics, consistent with quasi-periodic temporal structure with a characteristic period of approximately five minutes 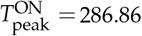, full width at half maximum (FWHM) = 16.92 s; 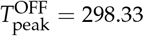 FWHM = 16.12 s).

In contrast, NEUTRAL-cells lacked consistent low-frequency spectral peaks, instead showing broad low-frequency components and prominent high-frequency power (*>* 10 Hz), reflecting regular, tonic firing. Although the prominence of the low-frequency peak varied across individual neurons, it was consistently observed at the population level for ON-and OFF-cells. These results reveal prominent low-frequency temporal structure in ON-and OFF-cell activity that was not consistently observed in the NEUTRAL-cell population.

We next summarized firing variability over the full recording period (Figure 3e). NEUTRAL-cells fired regularly and rapidly (Coefficient of Variation = 0.35 ± 0.26; mean Inter-Spike Interval = 0.17 ± 0.25 s, ∼ 10 Hz). In contrast, ON-and OFF-cells exhibited highly variable, burst-like firing patterns, with coefficients of variation well above 1 (CV_ON_ = 4.91 ± 4.32, CV_OFF_= 3.70 ± 3.09) and broad interspike interval (ISI) distributions (ISI_ON_= 1.25 ± 4.80 s, ISI_OFF_= 0.84 ± 1.36 s; Supplementary Table S1). Because relatively few neurons were recorded simultaneously in each animal, we did not attempt further within-class sub-clustering.

Together, these analyses indicate that ongoing ON-and OFF-cell activity contains structured low-frequency temporal organization, whereas NEUTRAL-cells maintain predominantly high-rate, non-rhythmic firing. However, frequency-domain measures such as PSD and summary statistics capture only average power and variability; they discard temporal phase information and cannot resolve how firing unfolds over time.

To address this, we next used Gaussian process (GP) modeling to characterize ongoing dynamics directly in the time domain. We implemented two complementary GP approaches: a purely periodic GP with a Gaussian likelihood to characterize the dominant low-frequency temporal structure, and a generalized Poisson GP with a SoftPlus link function to test whether these slow fluctuations were structured and predictable on a per-neuron basis.

### A periodic Gaussian process identifies dominant low-frequency temporal structure in ON-and OFF-cell activity

To identify the characteristic timescales underlying the slow fluctuations observed during ongoing activity, we applied a periodic GP model with a Gaussian likelihood to the smoothed firing rates of each neuron, scanning across 200 candidate periods (0.1–1500 s). This approach quantifies how well different candidate periods explain the observed firing pattern under the assumption that any repeating temporal structure is smooth and approximately periodic (Figure 4). ON-and OFF-cells showed clear minima in the negative log marginal likelihood (NLML) near 300 s, with additional subharmonic peaks at 600 and 900 s, consistent with a dominant low-frequency temporal structure having a characteristic period of approximately five minutes. In contrast, NEUTRAL-cells (with one exception) displayed flat NLML profiles with no consistent minima, without evidence of stable rhythmicity. The present analyses demonstrate statistical predictability rather than evidence for an intrinsic biological oscillator.

**Figure 4:**
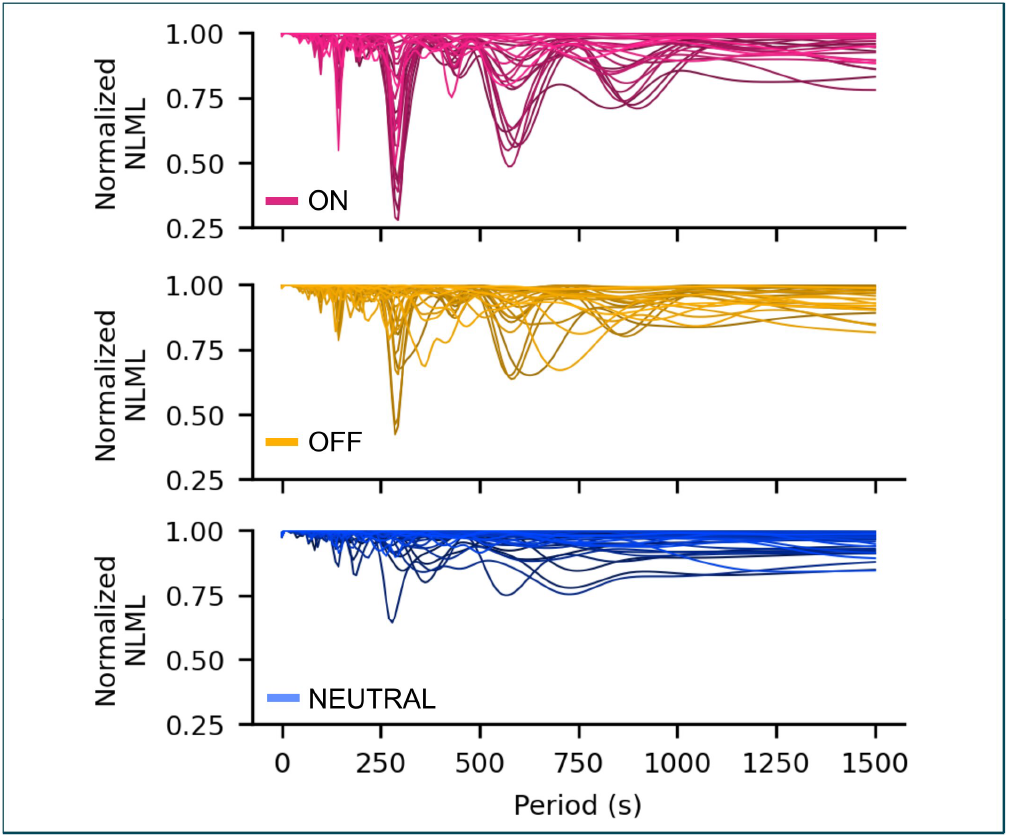
Period-length scans of the negative log marginal likelihood (NLML) from periodic Gaussian process fits to ongoing firing in ON-, OFF-, and NEUTRAL-cells. ON-and OFF-cells show distinct minima at ∼ 300 s and its harmonics, whereas NEUTRAL-cells exhibit flat profiles.

### A Poisson Gaussian process reveals predictable low-frequency dynamics in ongoing RVM firing

We next asked whether this low-frequency temporal structure was predictable, i.e. whether past firing reliably forecast future activity. To test this, we used a Poisson GP model with a SoftPlus link function, allowing the variance of firing rates to scale with their mean through a dispersion parameter (*α*). Each model was trained on all but the final 300 s of data and tasked with predicting the held-out test segment (Figure 5a). The fitted models (solid lines) accurately captured slow fluctuations during training and generalized into the test window (dotted lines) consistently for ON-cells, more variably for OFF-cells, and not for NEUTRAL-cells.

**Figure 5:**
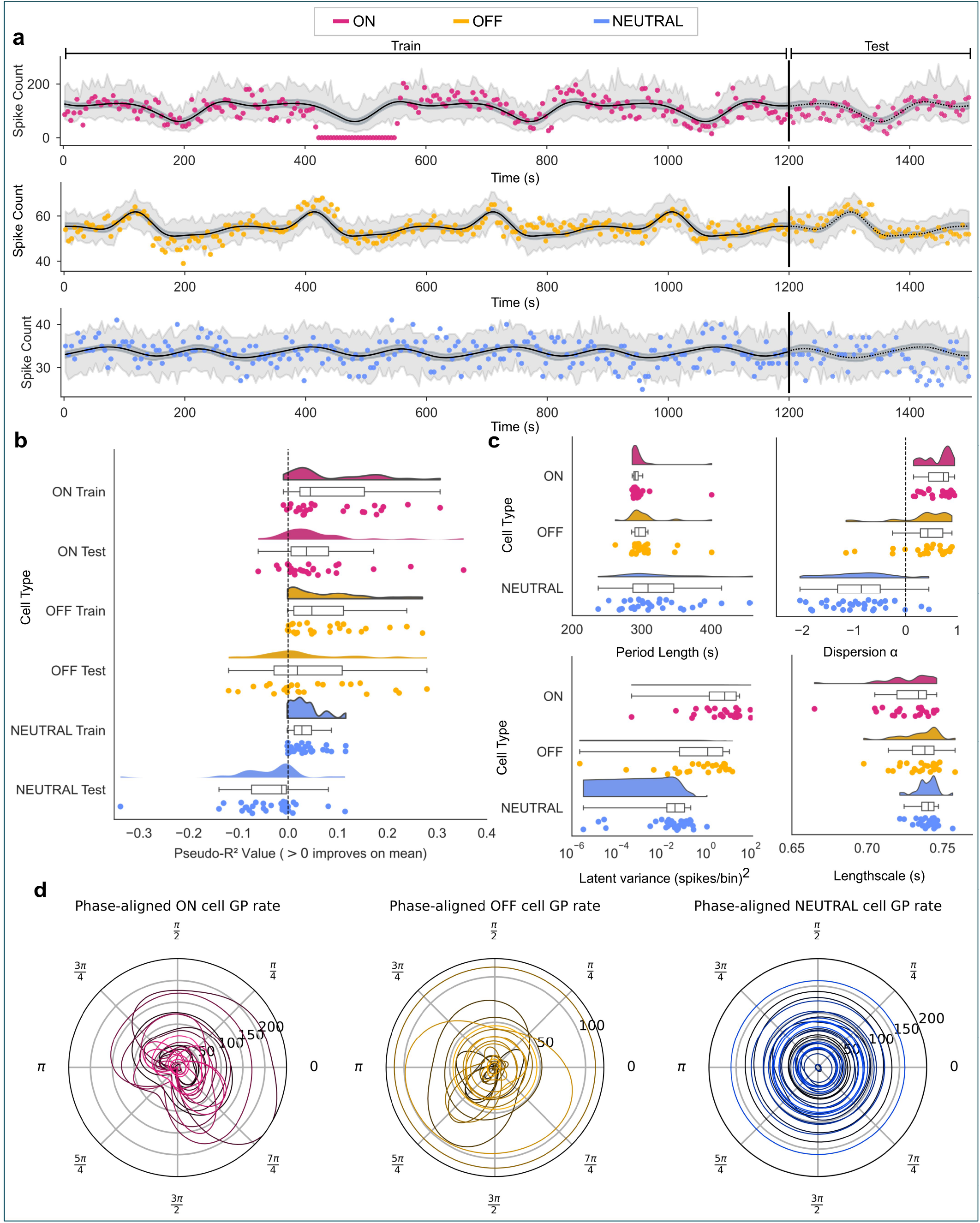
Periodic Gaussian process (GP) modeling of ongoing ON-, OFF-, and NEUTRAL-cell activity. **a**, Example fits for single ON-, OFF-, and NEUTRAL-cells using a periodic GP kernel. GP fits were robust to noise or occasional periods of silence in the training data (e.g. ON-cell trace, 410-550 s). Models were trained on all data except the final 300 s (training region: solid line) and tasked with predicting the held-out test segment (dotted line), without using test data during fitting. **b**, Pseudo-*R*^2^ values for training and test sets. ON-cells achieve significant predictive performance on test data, OFF-cells show weak predictability, while NEUTRAL-cells fail to generalize (test Pseudo-*R*^2^<0). **c**, Posterior hyperparameters across cell classes. ON-and OFF-cells exhibit positive dispersion parameters *α* and tightly clustered period estimates near 300 s, consistent with structured low-frequency temporal dynamics. NEUTRAL-cells show negative dispersion and less coherent period estimates, reflecting their regular, high-rate activity. **d**, Phase aligned GP rates adjusted for found period length showed unique signatures for each cell.

Training and test accuracies, quantified by pseudo-*R*^2^, were computed for each cell (Figure 5b). We performed two-sided T-tests to assess significance (*R*^2^ ≠ 0) using sign flipping permutations of the *R*^2^ value of all cells per animal, and the median *R*^2^ at each permutation (see Supplemental Figure S4 for distributions). This accounted for the potential correlation of ON-and OFF-cells recorded from the same animal. All three cell classes achieved significant training accuracy (*p*ON = 0.0051, *p*OFF = 0.0009, *p*NEUTRAL *<* 0.0001). Comparing *R*^2^ values using a similarly permuted and Bonferroni corrected Mann Whitney U test revealed no significant differences between training *R*^2^ for the three groups (see Supplemental Figure S6). This was expected, since even apparently aperiodic firing rates can contain some variance which can be captured by the GP. Therefore, we focused on predictive accuracy as a more stringent test of whether the low-frequency temporal structure captured by the periodic kernel generalized to unseen data.

By contrast, test predictability, quantified by pseudo-*R*^2^, dissociated the three populations (Figure 5b). ON-cells showed significant generalization to unseen data (*p* = 0.0091), with the large majority of cells exceeding baseline performance (i.e., >0). OFF-cells, as a population, did not show reliable predictive structure (*p* = 0.3145), reflecting a mixture of well-predicted and poorly predicted neurons. NEUTRAL-cells performed significantly below baseline (*p* = 0.0046), demonstrating a lack of periodicity. Group comparisons revealed significant differences between NEUTRAL-cells and both ON-and OFF-cells (*p <* 0.001 and *p* = 0.006, respectively), while ON-and OFF-cells did not differ significantly at test (*p* = 0.939). Kruskal–Wallis tests confirmed significant group differences at test time (*p <* 0.001) but not during training (*p* = 0.117). Thus, reproducible long-timescale predictability was most evident in ON-cells, while OFF-cells exhibited evidence of low-frequency temporal structure, although this was considerably more heterogeneous across neurons than in ON-cells.

Posterior hyperparameters (Figure 5c; Table 1) revealed further distinctions among cell classes. Estimated characteristic periods for both ON-and OFF-cells clustered near 300 s, corroborating the frequency-domain analyses. Although NEUTRAL-cells also displayed nominal period clustering near 300 s, their low latent variance and poor predictive performance indicate that this reflects weak constraint of the period parameter rather than meaningful periodic temporal structure.

**Table 1:** Posterior hyperparameters from the fitted latent GP model.

| Cell class | period $p$ (s) | dispersion $\alpha$ | outputscale $\sigma^2$ | lengthscale $l$ |
| --- | --- | --- | --- | --- |
| ON | $297 \pm 22$ | $0.63 \pm 0.25$ | $17 \pm 26$ | $0.73 \pm 0.018$ |
| OFF | $304 \pm 27$ | $0.36 \pm 0.53$ | $3.1 \pm 4$ | $0.74 \pm 0.013$ |
| NEUT. | $322 \pm 54$ | $-0.9 \pm 0.63$ | $0.1 \pm 0.2$ | $0.74 \pm 0.0068$ |

ON-and OFF-cells exhibited positive dispersion parameters (*α >* 0; *p*ON = 0.0011, *p*OFF = 0.0105), consistent with bursty, over-dispersed firing. In contrast, NEUTRAL-cells showed negative dispersion (*p*NEUTRAL *<* 0.001), indicating highly regular firing with variance below Poisson expectation. Latent GP variance further separated the classes: ON-cells showed the largest modulation amplitude, OFF-cells intermediate variance, and NEUTRAL-cells near-flat latent structure.

Finally, single-cycle phase-normalized reconstructions (Figure 5d) revealed reproducible phase-dependent firing profiles for ON-and OFF-cells, whereas NEUTRAL-cells showed near-constant phase profiles, consistent with an absence of structured modulation.

Together, these results demonstrate that slow low-frequency temporal structure is most consistently expressed and predictably organized in ON-cells, is present but heterogeneous in OFF-cells, and is largely absent in NEUTRAL-cells. The ability of the model to predict held-out activity indicates that ongoing firing in descending pain-modulatory neurons is not solely stochastic, but contains reproducible temporal organization operating on multi-minute timescales.

To compensate for the shorter recordings lengths of some NEUTRAL-cells in comparison to ON-and OFF-cell recordings, we refitted the Poisson GP model with equal amounts of training data (660 s) across all cells. The results, shown in Supplemental Figure S11, were as expected: the shorter training data resulted in decreased predictive performance for both ON-and OFF-cells; however, predictive performance was qualitatively similar in ordering to the full fit, with ON-cells predictions remaining the most consistent, and ON-and OFF-cells showing higher dispersion, latent variance, and a broader set of length scales to NEUTRAL-cells. We further tested our fitting method in simulated periodic and aperiodic cells (see Supplemental Figure S12), demonstrating the recovery of periodicity in 660 simulated recordings.

Although both ON-and OFF-cells exhibited similar characteristic timescales, the predictive structure was considerably more consistent across ON-cells than OFF-cells. Greater heterogeneity within the OFF-cell population may reflect differences in network coupling, modulation strength, or molecular composition, although these possibilities cannot be distinguished using the present data.

### Low-frequency coherence between RVM activity and heart rate

The RVM is implicated not only in nociceptive modulation but also in autonomic control, influencing cardiovascular and respiratory outputs ^34–39^. Given this dual role, we next asked whether the low-frequency temporal structure identified in RVM activity was also reflected in an autonomic variable, namely heart rate.

Previous studies using small samples of ON-and OFF-cells reported correlations between ON/OFF-cell firing and cardiovascular measures such as heart rate and blood pressure ^40,41^. Here, rather than testing instantaneous correlations, we quantified coherence between neuronal firing rates and simultaneously recorded heart rate. Coherence detects shared frequency-specific temporal structure and is robust to phase offsets and substantial uncorrelated variability.

The neuronal firing rates and simultaneously recorded heart rates (Figure 6a) were segmented, demeaned, and analyzed using multitaper coherence estimation (Figure 6b). Significant low-frequency coherence was observed in a subset of cells (5/32 NEUTRAL, 4/25 OFF, 8/25 ON; Figure 6c). Although the frequencies exhibiting significant coherence did not coincide exactly with the characteristic ∼ 300 s timescale identified in the neuronal analyses, the heart-rate power spectrum also contained prominent low-frequency components (Figure 6d), indicating that heart rate exhibits temporal structure over comparable timescales.

**Figure 6:**
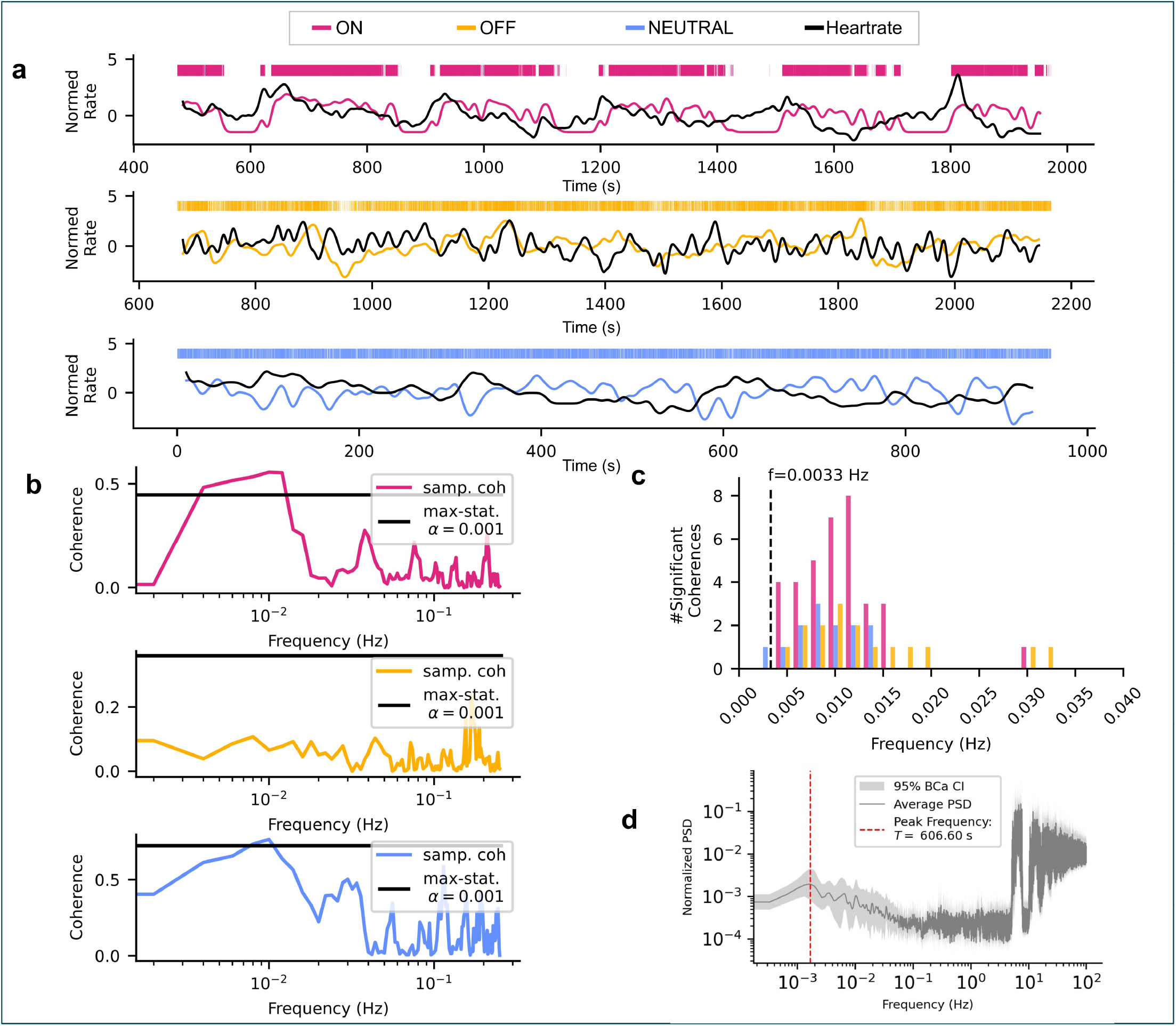
Analysis of heart rate coherence against cell firing rate. **a**, Example normed firing rates and heart rates for ON-, OFF-, and NEUTRAL-cells, showing both visibly coherent and incoherent activity. **b**, Coherence against frequency for the same cells in (**a**), showing the max-statistic significance level (black line). **c**, Distribution of significant coherence frequencies across all recorded neurons. **d**, Population-averaged heart-rate power spectrum. High-frequency peaks correspond to the cardiac rhythm, while additional low-frequency components indicate slower autonomic fluctuations. The lowest-frequency peak is highlighted in red.

These findings suggest that slow RVM dynamics can covary with autonomic activity in a subset of recordings. This shared temporal structure does not establish a causal interaction and could arise from common physiological influences acting on both signals.

## Discussion

Pain-regulatory circuits must coordinate rapid defensive responses with slower fluctuations in internal physiological state. Our results show that neurons in the rostral ventromedial medulla (RVM), a central hub of descending pain control, operate across distinct temporal scales. By combining neuronal recordings with probabilistic modeling, we find that identified ON-and OFF-cells exhibit both fast reflex-linked responses and slower, structured low-frequency dynamics during ongoing activity. Importantly, these features were not observed in NEUTRAL-cells recorded under the same anesthetic and physiological conditions, arguing against a uniform global influence affecting all RVM neurons similarly. Together, these findings indicate that descending pain-modulatory activity is organized across multiple timescales, although they do not by themselves establish the mechanistic origin or functional role of these dynamics.

Multi-timescale neural dynamics are well documented in cortical and neuromodulatory systems, where they are often attributed to large recurrent networks and global brainstate fluctuations ^42,43^. Our findings extend this principle to a brainstem circuit that directly regulates nociceptive transmission. In this framework, ON-and OFF-cells are not simple reciprocal antagonists. Rather, they contribute distinct and temporally distinct components of descending modulation—consistent with their different pharmacology and the fact that each population can be manipulated independently ^1^. Given the limited local connectivity among ON-and OFF-cells ^44^, the observed dynamics are unlikely to arise solely from local microcircuits and may instead reflect broader distributed feedback loops linking the RVM with spinal, midbrain, and hypothalamic control systems.

Bayesian modeling of withdrawal-aligned activity revealed that ON-and OFF-cells exhibit structured, multi-phase responses rather than simple bursts or pauses. Population activity was more tightly aligned to the withdrawal reflex than to the onset of noxious stimulation, confirming classical observations ^10^ while providing a quantitative description of the temporal structure of these responses. Importantly, the extrema of the two cell classes were temporally offset: ON-cell activity peaked 180 ms before the withdrawal, whereas OFF-cell suppression reached its nadir >2 seconds after. Thus, although the two populations are broadly anticorrelated, their dynamics are not simply mirrored but instead unfold in a temporally staggered manner.

We investigated both a sigmoidal and a Gaussian CDF function for the initial response of OFF-cells. Since we fitted our models to the summed population activity over animals and trials, we expect the slope of response function to be indicative of the general mechanism underlying the response. With this in mind, a sigmoidal decrease satisfies the logistic differential equation, in which there is recurrent inhibition (see Supplemental Gaussian CDF Fits). In contrast, a Gaussian CDF response suggests that OFF-cells cease firing independently around some response time *t*_0_. These two distributions are very difficult to distinguish between given the small timescale of the initial response, and the differences between the distributions are most pronounced in their tails, making the visual fits to data appear very similar. We observed that the sigmoidal function fit better to the response in terms of the ELPD; this is likely due to the heavier tails of the sigmoidal function fitting the noisy data better, and may change with a larger sample size or hierarchical model.

Analysis of the recovery phase revealed multiple timescales in both populations. ON-cell recovery was best captured by a double-exponential function, whereas OFF-cell recovery was best described by a combined linear and exponential trajectory. Neither population returned to its pre-stimulus firing rate within 100 s. This separation of fast and slow components suggests distinct biological processes underlying the initial response and the prolonged return toward baseline. The abrupt OFF-cell pause and sharp ON-cell peak are consistent with positive-feedback mechanisms that support rapid, precisely timed actions, whereas the slower recovery component aligns with the prolonged adjustments in nociceptive responsiveness observed behaviorally. Consistent with this, blocking the OFF-cell pause suppresses withdrawal ^45^, and attenuating the ON-cell burst reduces reflex magnitude ^13,46^. Moreover, the slow post-withdrawal drift toward baseline correlates with increased sensitivity to subsequent stimuli ^47^ and with altered ON/OFF-cell balance in persistent pain models (e.g., spinal nerve ligation, CFA inflammation ^48,49^). Together, these findings suggest that the initial response and its prolonged recovery serve complementary roles: coordinating the reflex itself while shaping a longer-lasting nociceptive state in the face of potential threat.

Beyond event-related responses, we identified structured low-frequency dynamics in ongoing firing, with a characteristic timescale of approximately five minutes. This temporal structure was evident in both the power spectrum and Gaussian process models and was reproducible across animals and cell classes, although it was not consistently observed in NEUTRAL-cells. The ability of Gaussian-process models to predict held-out activity indicates that these fluctuations are not simply stochastic variability but contain reproducible temporal organization. Importantly, statistical predictability alone does not establish that these dynamics arise from an intrinsic biological oscillator. Although ON-and OFF-cells exhibited similar characteristic timescales, the predictive structure was more consistent across ON-cells than OFF-cells. Greater heterogeneity within the OFF-cell population may reflect differences in network coupling, modulation strength, or molecular composition, although the present data cannot distinguish among these possibilities.

One possible interpretation is that ON-and OFF-cell activity reflects slowly varying network states that alter the balance of descending facilitation and inhibition. However, the present experiments cannot determine whether these dynamics originate within the RVM itself or arise from interactions with distributed circuits involved in autonomic, arousal, homeostatic, or other distributed brain networks. Rather than reflecting a generator within the RVM, the observed dynamics may instead represent coordinated activity within a broader pain-modulatory network.

An important unresolved question is the extent to which these slow fluctuations reflect dynamics within descending pain-modulatory circuits themselves versus coupling to broader physiological processes. The lightly anesthetized preparation used here was chosen because it enables stable recordings from physiologically identified RVM neurons while reducing the continuously varying sensory and behavioral influences that would otherwise complicate interpretation of their ongoing activity in awake animals, yet preserves the established relationships between ON-and OFF-cell activity and nociceptive processing. Although anesthesia likely influences global brain-state dynamics and may affect the amplitude, frequency, or coherence of the slow fluctuations described here, the absence of comparable temporal structure in NEUTRAL-cells argues against a uniform global effect acting similarly across all RVM neurons. At the same time, this observation cannot exclude systemic influences acting preferentially on ON-and OFF-cell circuitry. Determining which aspects of these dynamics generalize to awake animals, and how they relate to naturally occurring behavioral fluctuations, remains an important direction for future work.

Although we did not observe phase-dependent effects on withdrawal latency (Supplemental Figure S2), the experiment was not designed to test this question. Previous work demonstrating increased nociceptive sensitivity during periods of ON-cell dominance ^3,21^ suggests that these slowly varying network states may influence nociceptive responsiveness under experimental conditions specifically designed to address this issue.

Given the limited local connectivity among ON-and OFF-cells ^50^, and the lack of evidence for direct mono-synaptic connectivity between the populations ^44^, the low-frequency structure is unlikely to arise from intrinsic microcircuit loops and more plausibly reflects broader feedback interactions involving the spinal cord, other brainstem regions, hypothalamus, or forebrain ^3,6,18,22,51,52^. The consistency of the observed temporal structure across recordings suggests common modulatory influences within each functional population, although synchrony was not measured directly.

Because recordings were obtained from small pseudopopulations rather than large simultaneously recorded ensembles, we cannot determine the extent to which these fluctuations are synchronized across the broader ON-and OFF-cell populations. Future large-scale electrophysiological or imaging approaches will be required to establish how these dynamics are coordinated across the descending pain-modulatory network.

Furthermore, there is substantial molecular heterogeneity within ON-and OFF-cell populations ^6–9,53–55^. Because neurons were classified physiologically rather than molecularly, we cannot determine how the temporal dynamics described here relate to any identified molecular subclasses of ON-and OFF-cells. However, despite preventing a straightforward molecular classification of ON-and OFF-cells, such diversity may support the flexible functional organization observed here, enabling the integration of upstream control signals based on both internal (slow periodicity) and external (stimulus-evoked) variables. The RVM receives convergent non-nociceptive inputs ^56^ and participates in thermoregulatory and autonomic processes ^41,57^, further supporting a role as a multimodal hub integrating information across modalities and timescales.

Pain modulation is closely intertwined with autonomic and behavioral adjustments that support homeostasis ^23,58,59^. Prior work reports associations between ON/OFF-cell firing and heart rate or body temperature ^40,41^. Motivated by this, we examined heart rate as a potential covarying signal. Because both heart rate and RVM activity are shaped by many independent factors which tend to decorrelate signals, simple correlation is difficult to interpret; coherence provides a more targeted measure of shared structure at the timescale of interest (see Supplemental Coherence vs correlation). A subset of neurons across all three cell classes showed significant low-frequency coherence with heart rate, and heart-rate spectra exhibited low-frequency components spanning similar timescales. We interpret these findings cautiously. They suggest that a subset of RVM neurons participates in low-frequency dynamics that covary with autonomic activity, but they do not establish a causal relationship or a common generator.

Finally, the present dataset was predominantly male (77 males, 6 females), precluding meaningful assessment of sex differences. Although previous work suggests broadly comparable physiological properties of individual ON-and OFF-cells across sexes ^60^, whether the slow population dynamics described here generalize to females remains unknown.

In conclusion, our results demonstrate that descending pain-modulatory neurons operate across multiple timescales, from seconds-long reflex-related responses to minute-scale endogenous rhythms. This demonstrates that descending pain-modulatory activity contains structured dynamics even in the absence of noxious stimulation and places the RVM within a broader feedback loop balancing sensory processing, defensive behaviors, and internal physiological states. Characterizing this temporal organization provides a quantitative basis for understanding how brainstem pain-control circuits regulate nociception over time and may ultimately help inform future studies aimed at optimizing therapeutic engagement of descending pain pathways.

## Methods

### Experiments

All experiments followed the guidelines of the National Institutes of Health and the Committee for Research and Ethical Issues of the International Association for the Study of Pain, and were approved by the Institutional Animal Care and Use Committee at Oregon Health & Science University (OHSU).

Male and female (n = 77 and 6 respectively) Sprague-Dawley rats (Charles River; 191-348 g, mean ± SD: 264 ± 27 g) were deeply anesthetized using isoflurane (4-5%) and a catheter placed in the external jugular vein for subsequent infusion of the short-acting barbiturate methohexital. They were then transferred to a stereotactic frame and a small craniotomy made to gain access to the RVM. Body temperature was monitored and maintained at 36-37 °C with a heating pad, and heart rate was monitored using EKG. Mean ± SD heart rate was 391.1 ± 34.4 bpm, with mean range over the protocol 0.62 ± 12.0 bpm across all animals. Respiratory variables were not recorded and therefore could not be included in the present analyses.

After preparatory surgery was complete, the anesthetic plane was lowered such that a heat stimulus applied at 5-min intervals to the plantar surface of the hindpaw (radiant or contact heat delivered using a Peltier device) elicited a paw withdrawal. The stimulus consisted of a ramp from 36 to 53 °C over a period of 11 s. The stimulus was terminated when the animal withdrew the paw (mean ± SD paw withdrawal temperature: 49.0 ± 2.0 °C, latency: 8.7 ± 1.1 s). The paw surface was maintained at 35 °C between trials. Recordings were performed between August 2020 and August 2024.

Once the animal responded to the heat stimulus, methohexital was infused at a rate (37.8–77.5 mg/kg/hr, mean ± SD: 60.0 ± 7.6 mg/kg/hr) that maintained this lightly anesthetized state (as indicated by stable paw withdrawal response) while preventing spontaneous movement. Once established for a given animal, anesthetic flow rate was not changed for the remainder of the experiment. Accordingly, the ongoing recordings used for Gaussian-process analyses were obtained during periods of stable anesthetic infusion, without adjustment of methohexital rate.

These experiments used the well-validated lightly anesthetized Sprague-Dawley rat protocol in which much of the foundational data concerning physiology and function of RVM neurons was obtained. This “middle-out” strategy ^61^ has allowed direct connections between the activity and pharmacology of identified RVM neurons and altered nociceptive behavior. The protocol demonstrably spares the essential links between brainstem pain-modulating neurons and nociceptive transmission pathways. Although the focus here was on ongoing activity, precluding the repeated nociceptive testing needed to link neuronal activity to nociceptive threshold, previous work has demonstrated that ongoing activity of OFF-and ON-cells is correlated with nociceptive sensitivity ^21^ and that alterations in OFF-and ON-cell firing in response to pharmacological manipulation and in models of persistent pain states, stress, and sickness have behavioral relevance ^1,18^. Further, conclusions from work in lightly anesthetized rats have repeatedly been found to be congruent with behavioral observations by other groups in awake rats and mice ^6–8,55^ and with functional imaging evidence in humans ^2,62–64^. The lightly anesthetized model has thus established a circuit-level explanatory framework for behavioral findings obtained in several species in multiple laboratories.

The lightly anesthetized preparation was used here because the RVM is a deep brainstem structure and physiologically identified ON-and OFF-cells are relatively sparse. Prolonged single-unit recordings therefore require stable electrode– neuron contact, which can be readily disrupted by spontaneous movement. Methohexital light anesthesia suppressed spontaneous movement while preserving nocifensive withdrawal responses and the physiological response patterns required for ON-, OFF-, and NEUTRAL-cell classification ^28,65^. The experiment was performed in low ambient light conditions (< 10 lux). Extracellular single-unit recordings were acquired using a stainless-steel microelectrode (Frederick Haer & Co). Signals were amplified (10k) and band-pass filtered (400 Hz to 15 kHz, Neurolog, Digitimer) before analog-to-digital conversion at 32k samples/s for real-time spike detection and monitoring using Spike2 software (CED, Cambridge, UK). Correct waveform identification was verified on an individual spike basis at the conclusion of the experiment using Spike2 template matching and cluster analysis. EMG activity (to monitor withdrawal reflexes) was also recorded using Spike2. An RVM neuron or neurons was isolated and characterized as an ON-, OFF-, or NEUTRAL-cell based on changes in firing rate associated with withdrawal of the paw from a noxious mechanical stimulus applied to the hindpaw ^26,66^. ON-cells are defined by a period of activity beginning just prior to withdrawal from a noxious stimulus (or maintained firing if already active at the time of stimulus delivery). OFF-cells stop firing just prior to withdrawal (or remain silent if inactive). NEUTRAL-cells do not respond. We performed/analyzed data from two complementary protocols, each designed to capture different aspects of RVM activity during noxious stimulation or extended unstimulated periods. In the “evoked” protocol, noxious heat stimuli were delivered at 5 min intervals over a period of 16–18 min. A total of 104 cells were recorded in 53 animals: 35 OFF-cells, 45 ON-cells, and 24 NEUTRAL-cells. The “ongoing” protocol was designed specifically to examine ongoing activity in the absence of noxious stimulation. Cell activity was monitored for a period of 30 min following an initial noxious heat trial to classify the cell under study. Evoked responses were confirmed in at least two trials at the end of the experiment. Using this protocol, we recorded 62 cells in 30 animals: 25 OFF-cells, 25 ON-cells, and 12 NEUTRAL-cells. Together these protocols provided both noxious-evoked and prolonged recordings of ongoing, or “spontaneous” activity of ON-, OFF-, and NEUTRAL-cells, enabling systematic analysis of stimulus-driven dynamics and slow ongoing fluctuations. A detailed breakdown of cell numbers and animals used in each analysis is provided in Supplemental Table S11.

Recording sites in RVM were marked with an electrolytic lesion at the conclusion of the experiment. Animals were over-dosed with methohexital, and perfused transcardially with saline followed by 10% formalin. Brains were removed and post-fixed for 24 h in 10% formalin, then equilibrated for 24–72 h in 30% sucrose in PBS at 4 °C. Brains were sectioned at 40 to 60 µm, and recording sites were plotted (Supplementary Figure S13). The RVM was defined as the nucleus raphe magnus and adjacent reticular formation medial to the lateral boundary of the pyramids at the level of the facial nucleus.

### Data processing

After spike sorting and manual classification of cells, a window around each trial (-10 s to 100 s) was extracted relative to the alignment marker (heat onset or withdrawal behavior). Spikes within each window were aggregated across animals and trials to form the PSTHs for ON-and OFF-cells, providing a population-level view of stimulus-evoked activity. Trials were excluded from PSTH construction if the recording ended within the window or if the next event (heat onset or withdrawal) truncated the usable interval, resulting in a variable number of trials per PSTH.

For cells with available ongoing-activity recordings, we extracted a 1500 s segment (or 960 s for the shorter NEUTRAL-cell datasets) relative to the start of the first post–ongoing-activity trial. This ensured a minimum 230 s buffer after the initial trial, allowing transient evoked activity to dissipate before analyses of ongoing dynamics. Because noxious stimulation had no effect on NEUTRAL-cell firing rates, the full recording period from all neutral cells was used for GP modeling.

All analyses were conducted in Python v3.9. Instantaneous firing rates and basic statistics were computed using the Elephant package v1.1.1^67^. Gaussian processes were implemented using GPyTorch v1.13 and GPy v1.13.2^68,69^. Code interfaced with the Neo framework (v0.13) and the SonPy package (v1.9.5) to convert Spike2 (.smrx) files into Neo objects ^70,71^. Data was then standardized to the Neurodata Without Borders (NWB) standard ^72^. PyMC v5.12 was used for Bayesian regression ^73^.

We used Bayesian regression to model stimulation-evoked responses because the models contain relatively few parameters and the Bayesian framework provides full posterior uncertainty estimates—particularly useful for the switching timepoint. For ongoing activity, we employed maximumlikelihood type-II GP regression with a generalized Poisson likelihood. This approach was required because the long recording durations (> 1500 s), the nonstandard likelihood, and the multiple local minima induced by the periodic kernel made full Bayesian sampling computationally prohibitive.

### Piecewise Bayesian modeling of ON-and OFF-cells

We modeled the activity of a pseudo-population composed of cells and trials from multiple animals, rather than modeling each cell or trial individually in a multi-level model. The reasons for this were twofold: firstly, with only 3-4 trials per neuron, and roughly 2 neurons per animal, there was not enough data to estimate a full multi-level model. Secondly, as mentioned above, our primary aim was to capture population-level structure. Single-electrode recordings are essentially random samples from the population of ON-and OFF-cells, and we therefore assumed the emergence of population level effects to be due to exchangeability and uncorrelated noise averaging out across neurons.

We used piecewise Bayesian regression to fit the dynamics of ON-and OFF-cells during trials with noxious stimulation. We binned spikes using a 50 ms time bin, assuming that the total spike count for ON-or OFF-cells per bin 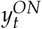 was Poisson distributed with a variable rate *λ*_*t*_, dependent on parameters *θ*, such that:

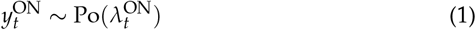

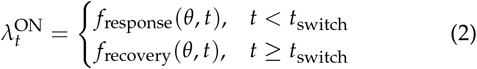

For ON-cells, the response phase was modeled as an exponential rise ^33^. We compared single-and double-timescale recovery functions. The double exponential model consisted of:

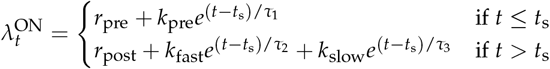

with continuity enforced by a corrected constraint:

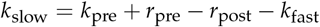

Priors were:

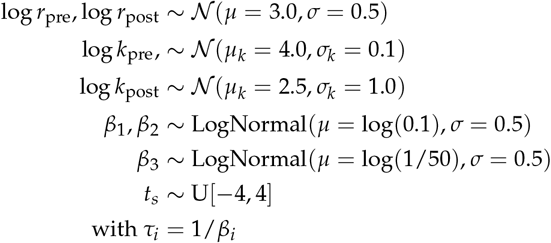

The single exponential model for ON-cells was identical except for removal of the *k*_fast_ term.

For OFF-cells we compared a sigmoidal and a gaussian CDF function as the response function ^33^ and compared a double timescale recovery with an exponential + linear recovery function. We constrained the sigmoid to pass through the point (*t*_*s*_, *u*_0_), joining the left and right side of the model. The general sigmoid has four parameters, slope *k*, midpoint *t*^∗^, lower asymptote *a* and upper asymptote *b*, such that:

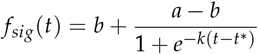

The Gaussian CDF had form of:

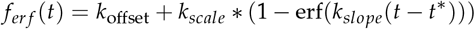

where erf refers to the Gauss error function. For both the Gaussian CDF and the sigmoid, we chose *t*^∗^ such that *f* (*t*_*s*_) = *u*_0_.

We fitted *a, b* and *k* directly; however, using a reparameterization of the sigmoid, we chose *t*^∗^ so that *f* (*t*_*s*_) = *u*_0_. Starting from the relation:

The full model consisted of:

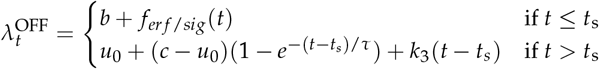

For the double exponential case, the right hand side of the model (*t > t*_*s*_) was replaced with:

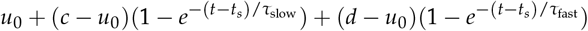

Priors were set as:

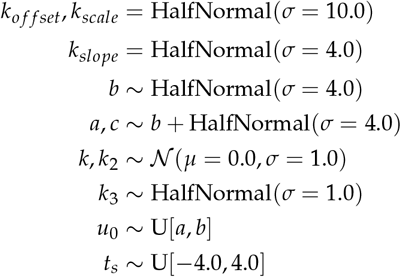

and additional priors for the double exponential OFF-cell recovery *d* and *k*_3_ were defined identically to *c* and *k*_2_ for the linear + exponential case, respectively. Markov chain Monte Carlo sampling using PyMC was used to sample from the posterior, using 1000 tuning steps, 4 chains, and 1000 samples per chain.

This Bayesian framework provided several advantages over prior methods ^33^: (i) it does not require predefining “burst” and “pause” times; (ii) it dissociates peak and nadir timing from withdrawal; (iii) it yields full posterior distributions for extrema and firing rates at any time; and (iv) it enables principled comparison of alternative timescale parameterizations for recovery dynamics.

### Gaussian process modeling

Firstly, as a preliminary period scan across potential timescales of ongoing firing, a GP with a periodic covariance function (see below) and Gaussian likelihood was applied to the smoothed firing rates of each neuron. We used 200 candidate periods (0.1-500 s) and a fixed lengthscale and variance of 1.0 and 5.0, respectively. A smoothing kernel and sampling period, both of width 2 seconds, were used to transform the spikes to a rate. The first and last 2 seconds of data were removed to avoid artifacts due to the smoothing process.

Secondly, slow structure in ongoing firing was quantified using latent GP models applied to binned spike counts (5 s bins). For each neuron, we fitted a sparse variational GP (200 inducing points) with a constant mean and a periodic covariance function:

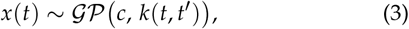

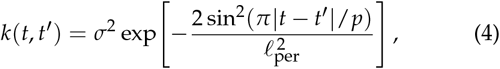

where *p* denotes the oscillation period and *l*_*p*er_ the periodic length scale. A narrow Gaussian prior *l*_per_ ∼ *N* (0.1, 0.1) was used to avoid over-smoothing in low-rate neurons. Kernel periods were initialized at 300 s, based on the period-scan analysis.

To map latent GP values to positive firing rates, a scaled Soft-Plus transform,

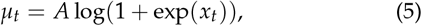

with *A* = 10 spikes/s was used instead of the exponential link to prevent numerical instabilities and speed up convergence across both low and high-rate regimes.

Spike count variability exhibited substantial over-and under-dispersion across cell classes. We evaluated a range of likelihoods (Poisson, Conway–Maxwell–Poisson), kernels (periodic, spectral mixture), and link functions. A generalized Poisson model ^74,75^ consistently provided stable fits and accommodated the broad dispersion range of ON-, OFF-, and NEUTRAL-cells. The generalized Poisson likelihood was parameterized by a parameter *θ*_*t*_ and dispersion parameter *α*:

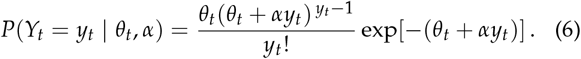

*θ*_*t*_ was related to the mean *µ*_*t*_ by:

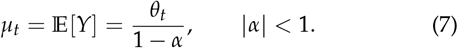

We placed a prior *α* ∼ *N* (0, 0.5) and computed *θ*_*t*_ = *µ*_*t*_ (1 − *α*) at each time point. A small contamination term (*p*_noise_ = 0.001) was added to account for rare outlier bins:

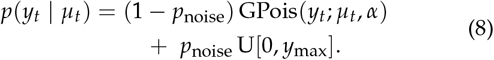

Finally, we confirmed the ability of our setup to find periodicity where present in the shorter 660 s training period of the NEUTRAL-cells, using a set of simulated periodic and aperiodic cells (see Supplemental Figure S12). We simulated cells using gaussian random noise on top of a mean firing rate, adding sinusoidal activity for the periodic cells, before clipping and rounding to integer non-negative spike counts. A 660 s training data duration is long enough to recover a 300 s oscillation, being more than double the period length. However, we acknowledge that the decreased amount of training data available for some NEUTRAL-cells may have, in principle, reduced the ability to estimate slow periodic structure present in those cells (see Discussion). Therefore, we also fitted our GP model to truncated data consisting of only the first 660 s of recordings, followed by a 300 s test period (see Supplemental S11).

### Coherence estimation

We first binned each spike train *x* and heart rate *y* using a 2 s bin size. We split the recording into 500 s segments with a 50% overlap, which gave each segment frequency resolution 1/*T* = 0.002 Hz. We demeaned each segment then used a multitaper spectral estimator with discrete prolate spheroidal sequences (DPSS) (standardized half bandwidth NW = 3.5, number of tapers *K* = 6) to calculate the coherence ^76^, using the formula:

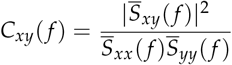

Segments were then individually phase-randomized to build a surrogate coherence distribution, and used a max-statistic approach to control the Family-Wise Error Rate (FWER) across all frequencies. The significance threshold was set at *α* = 0.001.

This combination of parameters gave a bandwidth = *B* = NW/*T* = 0.007 Hz, meaning that the frequency *f* = 0.0033 Hz (*T* = 5 minutes) found by the Gaussian Process was smeared across the low-frequency end of our coherence estimator. It was therefore not possible to determine whether the significant coherence found was due to oscillations at the same frequency as the GP period, due to the constraints of recording length and robust spectral estimation. Between individual cells, we observed some visual and spectral differences across all cell types between 0.01 Hz “fast” coherence and 0.0033 Hz “slow” coherence but these could not be quantified.

## Acknowledgments

FM is funded by a MRC Career Development Award (MR/T010614/1), a UKRI Advanced Pain Discovery Platform grant (MR/ W027593/1), and an EPSRC/MRC Programme Grant (UKRI1970). CA is funded by an EPSRC DTP (EP/W524633/1). The Heinricher laboratory is supported by grants from the National Institutes of Health (NS098660, NS120486) to MMH. CCdP is supported by a fellowship from the National Institute of Dental and Craniofacial Research (F31 DE030677).

## Declaration of interests

The authors declare no competing interests.

## Consent for publication

All authors gave their consent for publication.

## Data availability

All data related to this publication can be found on the DANDI data repository ^77^.

## Code availability

Code is available on the Zenodo code repository and on GitHub ^78^.

## Author Contributions

This study was performed via a collaboration between the Heinricher Lab at Oregon Health and Science University and the University of Cambridge. Carl Ashworth, Mary Heinricher and Flavia Mancini conceived the study. Melissa Martenson, Caitlynn De Preter, and Zhigang Shi performed the experiments. Carl Ashworth analyzed the data. Carl Ashworth, Mary Heinricher and Flavia Mancini wrote the manuscript. All authors reviewed the manuscript.

## Notes

### Competing Interest Statement

The authors have declared no competing interest.

### Summary of Updates

An updated abstract, title, and additional textual qualifications + supplementary analyses based on truncated/simulated data, following peer review by eLife.

