## Supplementary Information for "Multi-timescale dynamics in descending pain modulation"

#### Spike train power spectra

A spike train can be represented by a sum of delta functions $s(t) = \sum_{i=0}^{i=N} \delta(t - t_i)$. Therefore, the spectrum for a spike train can be directly computed as a sum over complex exponentials at each spike location, which can be evaluated at any frequency. However, due to the number of spikes and the computational benefits of using the FFT algorithm to compute the PSD, it is conventional to first bin the spikes with a fine bin width (e.g. 5 ms). The resulting sequence, which tends to a binary sequence as the bin width becomes small relative to the average spacing between spikes, is then used to compute the PSD using conventional FFT methods. This then restricts the frequencies to a discrete grid of frequencies up to the Nyquist frequency.

We used a 5ms bin width, giving a maximum realizable frequency of $\frac{1}{2f_s} = 100$ Hz. The lowest realizable frequency is given by $\frac{1}{T}$, where $T$ is the total recording time. For the ON- and OFF-cells this was $\frac{1}{1500} = 0.00067$ Hz, and for the NEUTRAL-cells this was $1/960 = 0.001$ Hz and $\frac{1}{3000} = 0.00033$ Hz. For comparability of the combined frequency plot for the NEUTRAL-cells the sequences were zero-padded to 100 times the length of the shorter recording.

Hierarchical bootstrapping at the rat level was used to generate confidence intervals for the average PSD. $N$ rats (equal to the sample number of rats) were resampled at each bootstrap iteration, using all spike trains from that rat. Per bootstrap iteration, the highest magnitude peak was then taken, and the FWHM computed. 1000 bootstrap iterations were used. Hierarchical bootstrapping helps to combat the decreased variance in the bootstrapped confidence intervals caused by correlation between spike trains measured from the same rat. Examples of this correlation are the higher likelihood of similar spectral peaks being observed, and the shadowing effect of a small number of correlated samples.

In addition to the main spectral peaks for the ON- and OFF-cells, the bootstrapped spectra also show the characteristic plateau at higher frequencies (Figure 3). This is characteristic of the PSD of a spike train; for example, Poisson neurons have a flat power spectrum corresponding to the average firing rate, with a single delta function at 0. Interestingly, there are several peaks in the NEUTRAL-cell PSD above 10 Hz. These, along with the low CV of the NEUTRAL-cells, show that they are more regular than Poisson firing neurons (CV = 1).

Finally, bootstrapping was used to compute the power spectrum for the heart rate in the same manner as for the cells. Here, there was no need to adjust for cluster sizes since each heart rate was only sampled once.

#### Technicalities of GP fitting

Here we discuss the benefits of using the raw binned spikes for GP fitting instead of the Gaussian kernel smoothed spike trains.

One approach to the modeling the firing rate is to convolve the spike train with a Gaussian kernel at each time point:

$$x(t) = \int d\tau \phi(\tau) s(t - \tau)$$

From this we obtain the average intensity of the single spike train. We then consider this intensity $x(t)$ to be a draw from a Gaussian process ($x \sim \mathcal{GP}$). By using a Periodic kernel, this model was able to fit well to ON- and OFF-cells, predicting their periodic behavior. However, the model leaves open the possibility of negative firing rates, and makes several false assumptions. Firstly, the model allows negative firing rates to be modeled by the GP. Secondly, since we have a constant density of points, the variance of the GP under this model does not decrease with the magnitude of $x(t)$, whereas for a more plausible model of spike trains, the lower the spike density, the lower the variance of the firing rate estimate. Finally, the GP assumes that the joint density of all points is Gaussian distributed. This assumption presents issues with overfitting in the case of spike trains, because we constructed the intensity estimate $x(t)$ using a Gaussian kernel. Therefore, there is a particular length scale at which data is well-fitted by an RBF kernel, leading to overfitting. This is not an issue when using a purely periodic kernel to model the firing rate; however, it would become an issue when modeling the trends in the firing rate with a periodic * RBF kernel, as the RBF kernel easily overfits to the data. One alternative for future study would be to use two separate RBF kernels: the first additive, modeling the short scale covariance in the firing rate induced by the Gaussian kernel, and the second kernel multiplying the periodic kernel.

However, convolving the firing rate estimate first with a kernel misses the fundamental issue with this method: spike trains are inherently noisy, and if we use a kernel to produce a continuous signal before fitting the spikes, we do not gain any new information, and remove the noise which prevents the GP from overfitting.

Therefore, a better approach is to bin the spike train. We make the assumption that the spike counts in each bin are independent, so that individual bin counts are dependent only on the value of the GP at that time point. From there, we assume that the GP is the underlying rate function for a discrete distribution (Poisson or otherwise) from which spikes are drawn. This approach deals with the fact that spike counts are integers whereas the GP models continuous functions.

Overall, the fit itself is less important than evaluating the prediction of the GP on the unseen firing rate data, as due to the noise, there will always be non-zero power at all frequencies in the training data, to which the GP can fit.

#### Ongoing spike train statistics

| Cell Type | CV (mean ± std) | ISI (s) (mean ± std) |
|---|---|---|
| Neutral | 0.348 ± 0.264 | 0.165 ± 0.250 |
| OFF | 3.699 ± 3.087 | 0.835 ± 1.357 |
| ON | 4.909 ± 4.319 | 1.248 ± 4.799 |

Table S1: The aggregated mean CV and ISI values of each of the three ongoing cell populations.

#### Bayesian Modeling Percentage Peak and Nadir Sampled Statistics

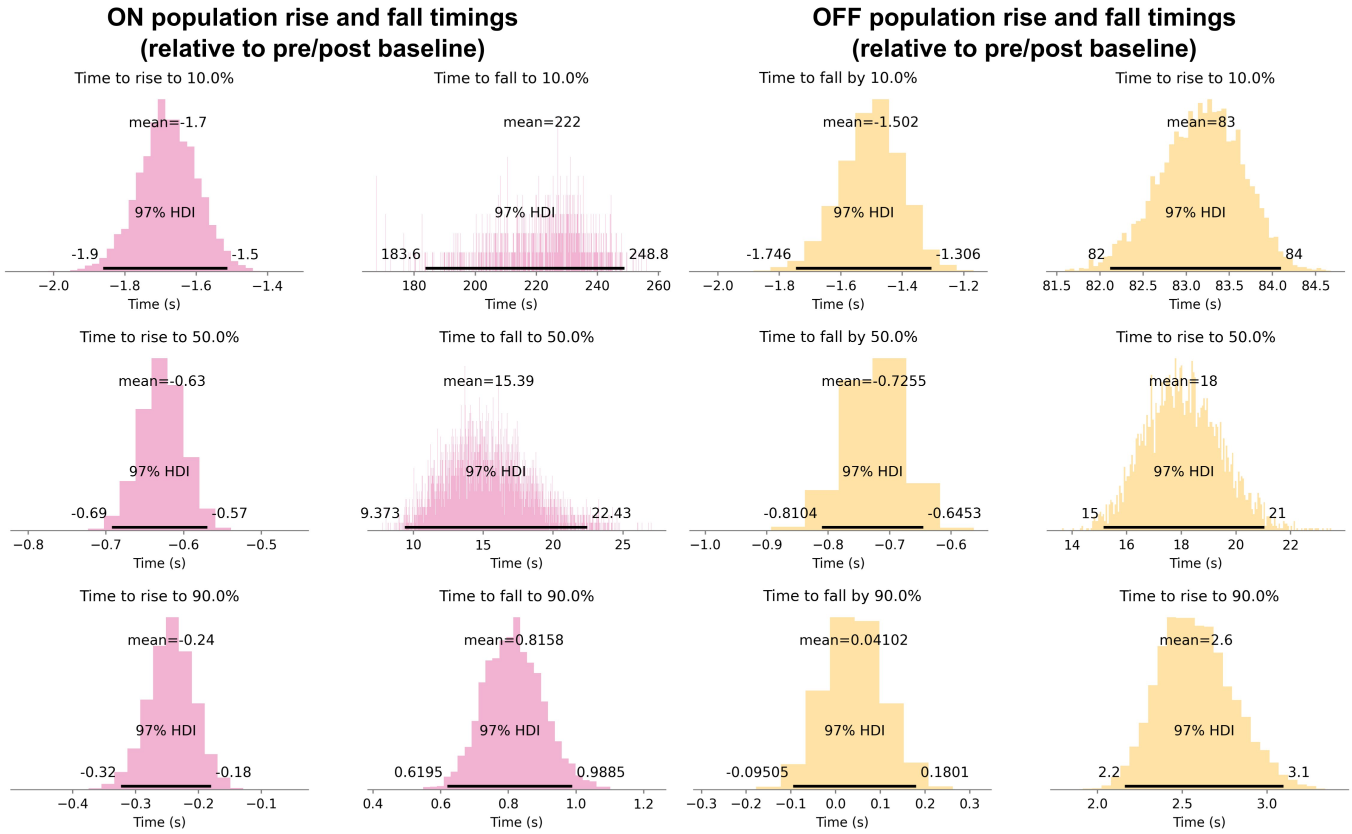


Figure S1: Posterior Distributions for the time to 10, 50 and 90% of peak for ON- cells, and 10,50 and 90% of the nadir for OFF-cells, with HDIs.

| Percentile | ON cells | | OFF cells | |
|---|---|---|---|---|
| | Rise(s) | Fall(s) | Fall(s) | Rise(s) |
| 10% | $-1.7$ [$-1.9$, $-1.5$] | 222 [183.6, 248.8] | $-1.502$ [$-1.746$, $-1.306$] | 83 [82, 84] |
| 50% | $-0.63$ [$-0.69$, $-0.57$] | 15.39 [9.373, 22.43] | $-0.7255$ [$-0.8104$, $-0.6453$] | 18 [15, 21] |
| 90% | $-0.24$ [$-0.32$, $-0.18$] | 0.8158 [0.6195, 0.9885] | 0.04102 [$-0.09505$, 0.1801] | 2.6 [2.2, 3.1] |

Table S2: Mean and 97% HDI for rise and fall times in ON and OFF cells at 10%, 50%, and 90% percentiles. Rises and falls for ON-cells are measured from *base to peak*. Rises and falls for OFF-cells are measured from *base to nadir*.

#### Ongoing activity phase did not affect withdrawal latency

Given that ON- and OFF-cells exhibited slow, quasi-periodic fluctuations in their ongoing activity, we next asked whether the phase of these rhythms influenced behavioral responsiveness to noxious stimuli. In principle, if RVM activity modulates nociceptive gain over time, withdrawal latency might vary systematically with the predicted phase of ON- and OFF-cell activity at stimulus onset.

However, the present experiments were not designed to investigate the relationship between cell dynamics and nociceptive sensitivity. Heat stimuli were not delivered at fixed intervals but rather during periods chosen to facilitate cell classification (OFF-cells active, ON-cells silent). Moreover, contact (Peltier) and radiant heat were used in different experiments, and the number of trials for each cell was small.

Despite these constraints, we explored whether withdrawal latency correlated with the phase or predicted firing rate derived from the periodic GP model fit to each neuron's ongoing activity.

As shown in Supplementary Figure S2d,e,f, there was no systematic relationship between normalized withdrawal latency and either predicted firing rate or oscillatory phase at stimulus onset. This suggests that, under the present recording and stimulation conditions, the slow RVM oscillations did not measurably modulate reflex latency. Nevertheless, prior studies designed explicitly for this purpose have reported that nociceptive sensitivity is increased during periods of ON-cell dominance[21,47], consistent with a potential behavioral impact of these network states.

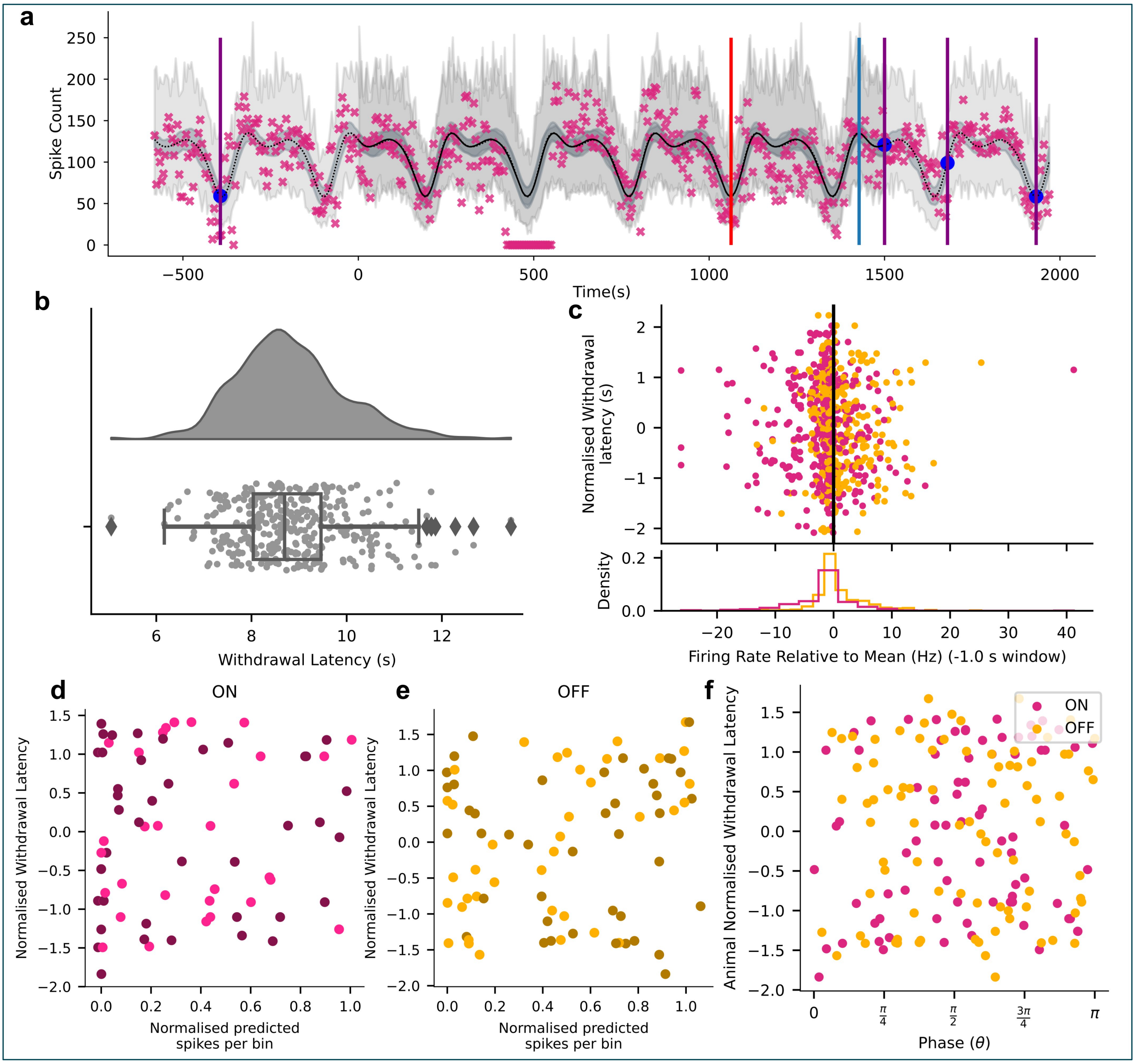


Figure S2: **a**, Example ON-cell GP fit from 0 to 1500 seconds, with extrapolated firing rate to trial times, indicated with dark blue circles. Minimum firing rate (red) and maximum firing rate (turquoise) were used to calculate the GP phase at each trial time. **b**, The distribution of withdrawal latencies across all trials and animals. **c**, The mean firing rate relative to mean of each cell before a trial, and the corresponding withdrawal latency. There was no visible effect of pre-heat firing rate on withdrawal latency, although a slight separation between ON (pink) and OFF cells (yellow) along the x-axis can be seen, potentially due to the difficulty of observing an OFF-cell if it was not already firing, and vice-versa for ON-cells. **d**, The spike count per bin predicted by the GP, normalized between the max and minimum spike count per GP fit, showed no visible correlation with withdrawal latency normalized per cell for ON-cells (**e**) or OFF-cells (**f**). The predicted GP phase showed no visible correlation with normalized withdrawal latency.

#### Neutral cell PSTHS

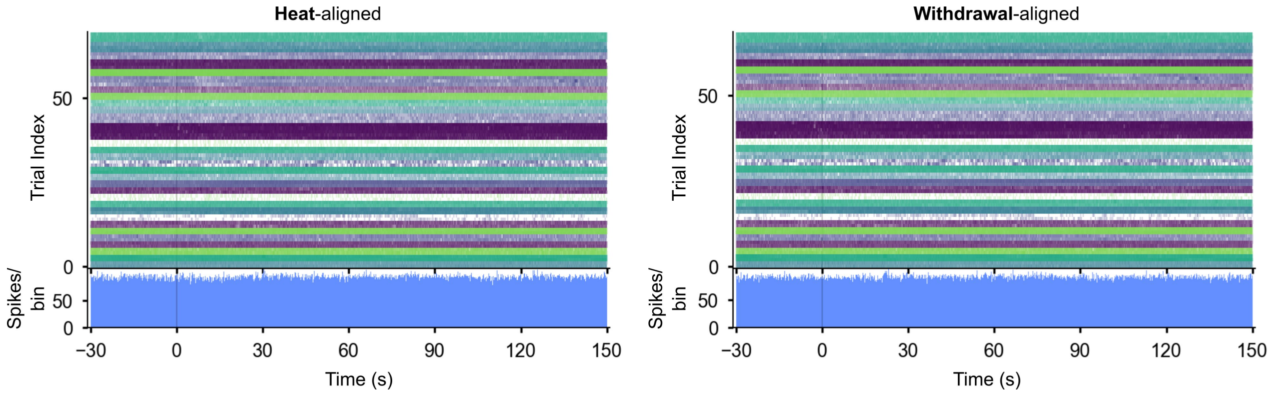


Figure S3: NEUTRAL-cells PSTHs showed no consistent modulatory effect on NEUTRAL-cells by the noxious heat or withdrawal response, nor at any time over the duration of the trial.

#### Bayesian trial model posteriors

Throughout this section, $r_{hat} > 1.0$ indicates that the model had difficulty fitting, i.e. that the samples were either bimodal (in the case of the double exponential OFF-cell model) or that the parametric form was likely not fully correct for the data (e.g. the heat aligned ON-cell). *posterior_rate_diff* is the difference between the pre-stimulus baseline and the post-behavior recovery baseline (for the double exponential models).

##### ON-cells

| | mean | sd | hdi_3% | hdi_97% | mcse_mean | mcse_sd | ess_bulk | ess_tail | r_hat |
|---|---|---|---|---|---|---|---|---|---|
| r_post | 16.002 | 5.129 | 6.475 | 25.429 | 0.112 | 0.081 | 2148.000 | 2205.000 | 1.000 |
| r_pre | 30.685 | 0.473 | 29.822 | 31.574 | 0.009 | 0.006 | 3085.000 | 2544.000 | 1.000 |
| k_post | 50.007 | 2.232 | 45.764 | 54.248 | 0.054 | 0.039 | 1691.000 | 2041.000 | 1.000 |
| k_slow | 64.386 | 4.881 | 54.621 | 72.667 | 0.104 | 0.074 | 2234.000 | 2297.000 | 1.000 |
| tau_post_slow | 164.223 | 18.883 | 129.571 | 199.809 | 0.417 | 0.295 | 2089.000 | 2168.000 | 1.000 |
| t_switch | -0.184 | 0.039 | -0.257 | -0.108 | 0.001 | 0.001 | 2393.000 | 2311.000 | 1.000 |
| k_pre | 99.711 | 2.318 | 95.487 | 104.250 | 0.058 | 0.041 | 1586.000 | 1807.000 | 1.000 |
| tau_post_fast | 4.157 | 0.309 | 3.582 | 4.722 | 0.007 | 0.005 | 1905.000 | 2254.000 | 1.000 |
| tau_pre | 0.653 | 0.046 | 0.570 | 0.739 | 0.001 | 0.001 | 2694.000 | 2229.000 | 1.000 |
| posterior_rate_diff | 14.682 | 5.159 | 4.901 | 23.911 | 0.113 | 0.080 | 2131.000 | 2143.000 | 1.000 |

Table S3: ON-cell withdrawal-aligned model posterior, using the double exponential model.

| | mean | sd | hdi_3% | hdi_97% | mcse_mean | mcse_sd | ess_bulk | ess_tail | r_hat |
|---|---|---|---|---|---|---|---|---|---|
| r_post | 53.642 | 0.589 | 52.527 | 54.751 | 0.014 | 0.010 | 1864.000 | 2493.000 | 1.000 |
| r_pre | 30.638 | 0.467 | 29.740 | 31.514 | 0.010 | 0.007 | 2401.000 | 2451.000 | 1.000 |
| t_switch | -0.382 | 0.040 | -0.456 | -0.308 | 0.001 | 0.001 | 2474.000 | 2541.000 | 1.000 |
| k_pre | 73.046 | 1.141 | 70.978 | 75.224 | 0.025 | 0.018 | 1995.000 | 2656.000 | 1.000 |
| k_post | 50.042 | 0.830 | 48.499 | 51.634 | 0.013 | 0.010 | 3828.000 | 3019.000 | 1.000 |
| tau_post | 26.095 | 1.398 | 23.530 | 28.810 | 0.035 | 0.025 | 1660.000 | 2223.000 | 1.000 |
| tau_pre | 0.659 | 0.053 | 0.566 | 0.761 | 0.001 | 0.001 | 2502.000 | 2214.000 | 1.000 |

Table S4: ON-cell withdrawal-aligned model posterior, using the single exponential model.

| | mean | sd | hdi_3% | hdi_97% | mcse_mean | mcse_sd | ess_bulk | ess_tail | r_hat |
|---|---|---|---|---|---|---|---|---|---|
| r_post | 12.347 | 4.233 | 4.892 | 20.210 | 0.104 | 0.078 | 1699.000 | 1699.000 | 1.000 |
| r_pre | 30.157 | 0.335 | 29.530 | 30.784 | 0.006 | 0.004 | 3003.000 | 2691.000 | 1.000 |
| k_post | 33.527 | 1.931 | 29.894 | 37.262 | 0.048 | 0.034 | 1604.000 | 1730.000 | 1.000 |
| k_slow | 70.232 | 4.131 | 62.556 | 77.556 | 0.098 | 0.069 | 1826.000 | 2210.000 | 1.000 |
| tau_post_slow | 5.115 | 0.437 | 4.325 | 5.963 | 0.010 | 0.007 | 2040.000 | 2510.000 | 1.000 |
| t_switch | 8.694 | 0.081 | 8.542 | 8.844 | 0.002 | 0.001 | 1817.000 | 1820.000 | 1.000 |
| k_pre | 85.949 | 1.888 | 82.645 | 89.676 | 0.049 | 0.035 | 1496.000 | 1770.000 | 1.000 |
| tau_post_fast | 170.341 | 14.593 | 141.835 | 195.972 | 0.361 | 0.255 | 1671.000 | 1530.000 | 1.000 |
| tau_pre | 1.278 | 0.079 | 1.129 | 1.426 | 0.002 | 0.001 | 2318.000 | 2503.000 | 1.000 |

Table S5: ON-cell heat-aligned model posterior, using the double exponential model.

**OFF-cells**

| | mean | sd | hdi_3% | hdi_97% | mcse_mean | mcse_sd | ess_bulk | ess_tail | r_hat |
|---|---|---|---|---|---|---|---|---|---|
| a | 46.355 | 0.376 | 45.687 | 47.083 | 0.006 | 0.004 | 4525.000 | 3151.000 | 1.000 |
| b | 7.559 | 0.574 | 6.556 | 8.680 | 0.018 | 0.012 | 1078.000 | 1375.000 | 1.000 |
| c | 20.358 | 0.324 | 19.775 | 20.974 | 0.007 | 0.005 | 2455.000 | 2657.000 | 1.000 |
| k | 1.747 | 0.152 | 1.476 | 2.041 | 0.004 | 0.003 | 1619.000 | 2191.000 | 1.000 |
| k2 | 0.286 | 0.040 | 0.213 | 0.359 | 0.001 | 0.001 | 1868.000 | 1992.000 | 1.000 |
| tau_post | 3.566 | 0.488 | 2.667 | 4.486 | 0.011 | 0.008 | 1868.000 | 1992.000 | 1.000 |
| k3 | 0.177 | 0.006 | 0.165 | 0.188 | 0.000 | 0.000 | 2300.000 | 2330.000 | 1.000 |
| t_switch | 11.718 | 0.287 | 11.198 | 12.269 | 0.008 | 0.005 | 1395.000 | 2063.000 | 1.000 |
| u0 | 7.697 | 0.504 | 6.781 | 8.640 | 0.015 | 0.010 | 1178.000 | 1662.000 | 1.000 |

Table S6: OFF-cell heat-aligned model posterior, using the exponential+linear model. $\tau_{post} = 1/k_2$.

| | mean | sd | hdi_3% | hdi_97% | mcse_mean | mcse_sd | ess_bulk | ess_tail | r_hat |
|---|---|---|---|---|---|---|---|---|---|
| a | 46.500 | 0.544 | 45.472 | 47.501 | 0.008 | 0.006 | 4484.000 | 2959.000 | 1.000 |
| b | 5.683 | 0.581 | 4.582 | 6.740 | 0.016 | 0.012 | 1272.000 | 1746.000 | 1.000 |
| c | 18.803 | 0.349 | 18.125 | 19.439 | 0.008 | 0.005 | 2143.000 | 2392.000 | 1.000 |
| k | 2.852 | 0.282 | 2.333 | 3.392 | 0.006 | 0.005 | 1863.000 | 2404.000 | 1.000 |
| k2 | 0.213 | 0.026 | 0.163 | 0.260 | 0.001 | 0.000 | 1960.000 | 2223.000 | 1.000 |
| tau_post | 4.760 | 0.577 | 3.711 | 5.861 | 0.013 | 0.009 | 1960.000 | 2223.000 | 1.000 |
| k3 | 0.186 | 0.006 | 0.174 | 0.196 | 0.000 | 0.000 | 1928.000 | 1902.000 | 1.000 |
| t_switch | 1.346 | 0.249 | 0.904 | 1.832 | 0.007 | 0.005 | 1313.000 | 1658.000 | 1.000 |
| u0 | 5.839 | 0.496 | 4.947 | 6.781 | 0.013 | 0.009 | 1478.000 | 1928.000 | 1.000 |

Table S7: OFF-cell withdrawal-aligned model posterior, using the exponential+linear model. $\tau_{post} = 1/k_2$.

| | mean | sd | hdi_3% | hdi_97% | mcse_mean | mcse_sd | ess_bulk | ess_tail | r_hat |
|---|---|---|---|---|---|---|---|---|---|
| a | 46.762 | 0.538 | 45.801 | 47.848 | 0.008 | 0.006 | 4566.000 | 2868.000 | 1.000 |
| b | 5.303 | 0.654 | 4.140 | 6.573 | 0.020 | 0.014 | 1114.000 | 1592.000 | 1.000 |
| c | 15.802 | 0.535 | 14.765 | 16.789 | 0.011 | 0.008 | 2238.000 | 2892.000 | 1.000 |
| d | 33.068 | 1.751 | 30.135 | 36.547 | 0.043 | 0.030 | 1749.000 | 1821.000 | 1.000 |
| k | 2.757 | 0.281 | 2.245 | 3.289 | 0.007 | 0.005 | 1750.000 | 2316.000 | 1.000 |
| k2 | 0.353 | 0.070 | 0.237 | 0.486 | 0.002 | 0.001 | 1908.000 | 2289.000 | 1.000 |
| k3 | 0.013 | 0.002 | 0.010 | 0.016 | 0.000 | 0.000 | 1495.000 | 1712.000 | 1.000 |
| tau_post_fast | 2.942 | 0.557 | 1.881 | 3.959 | 0.013 | 0.009 | 1908.000 | 2289.000 | 1.000 |
| tau_post_slow | 79.021 | 10.422 | 60.994 | 99.559 | 0.274 | 0.196 | 1495.000 | 1712.000 | 1.000 |
| t_switch | 1.488 | 0.300 | 0.976 | 2.037 | 0.008 | 0.006 | 1399.000 | 2105.000 | 1.000 |
| u0 | 5.455 | 0.545 | 4.450 | 6.480 | 0.016 | 0.011 | 1213.000 | 1910.000 | 1.000 |
| posterior_rate_diff | 3.346 | 2.035 | -0.401 | 7.207 | 0.052 | 0.037 | 1604.000 | 1864.000 | 1.000 |

Table S8: OFF-cell withdrawal-aligned model posterior, using the double exponential model.

| | mean | sd | hdi_3% | hdi_97% | mcse_mean | mcse_sd | ess_bulk | ess_tail | r_hat |
|---|---|---|---|---|---|---|---|---|---|
| erfscale | 21.296 | 0.440 | 20.437 | 22.098 | 0.013 | 0.009 | 1193.000 | 1235.000 | 1.000 |
| erfoffset | 4.666 | 0.666 | 3.448 | 5.922 | 0.021 | 0.015 | 997.000 | 921.000 | 1.000 |
| erfslope | 1.048 | 0.098 | 0.861 | 1.228 | 0.003 | 0.002 | 1456.000 | 1368.000 | 1.000 |
| c | 15.783 | 0.553 | 14.757 | 16.846 | 0.014 | 0.010 | 1626.000 | 2243.000 | 1.000 |
| d | 32.536 | 1.650 | 29.693 | 35.583 | 0.044 | 0.031 | 1526.000 | 1872.000 | 1.000 |
| k2 | 0.319 | 0.054 | 0.231 | 0.428 | 0.002 | 0.001 | 1175.000 | 751.000 | 1.000 |
| k3 | 0.013 | 0.002 | 0.010 | 0.016 | 0.000 | 0.000 | 1210.000 | 1533.000 | 1.000 |
| tau_post_fast | 3.221 | 0.538 | 2.312 | 4.265 | 0.018 | 0.015 | 1175.000 | 751.000 | 1.000 |
| tau_post_slow | 79.543 | 10.008 | 62.497 | 98.951 | 0.304 | 0.224 | 1210.000 | 1533.000 | 1.000 |
| t_switch | 1.170 | 0.243 | 0.721 | 1.656 | 0.009 | 0.006 | 837.000 | 669.000 | 1.000 |
| u0 | 4.861 | 0.529 | 3.928 | 5.890 | 0.015 | 0.011 | 1244.000 | 1884.000 | 1.000 |

Table S9: OFF-cell withdrawal-aligned model posterior, using the Gaussian CDF model with a double exponential recovery.

| | mean | sd | hdi_3% | hdi_97% | mcse_mean | mcse_sd | ess_bulk | ess_tail | r_hat |
|---|---|---|---|---|---|---|---|---|---|
| erfscale | 21.283 | 0.462 | 20.390 | 22.115 | 0.021 | 0.015 | 501.000 | 489.000 | 1.000 |
| erfoffset | 4.664 | 0.693 | 3.281 | 5.798 | 0.033 | 0.024 | 447.000 | 265.000 | 1.010 |
| erfslope | 1.075 | 0.105 | 0.896 | 1.279 | 0.003 | 0.002 | 1122.000 | 1378.000 | 1.000 |
| c | 18.825 | 0.353 | 18.146 | 19.463 | 0.009 | 0.007 | 1420.000 | 2101.000 | 1.000 |
| k2 | 0.204 | 0.023 | 0.163 | 0.246 | 0.001 | 0.000 | 1289.000 | 1004.000 | 1.000 |
| k3 | 0.184 | 0.006 | 0.172 | 0.194 | 0.000 | 0.000 | 1465.000 | 1884.000 | 1.000 |
| tau_post_fast | 4.957 | 0.546 | 4.027 | 6.099 | 0.016 | 0.011 | 1289.000 | 1004.000 | 1.000 |
| t_switch | 0.978 | 0.206 | 0.591 | 1.350 | 0.008 | 0.006 | 676.000 | 458.000 | 1.000 |
| u0 | 4.980 | 0.494 | 4.036 | 5.860 | 0.017 | 0.012 | 800.000 | 1136.000 | 1.010 |

Table S10: OFF-cell withdrawal-aligned model posterior, using the Gaussian CDF model with a linear + exponential recovery.

#### Permutation tests

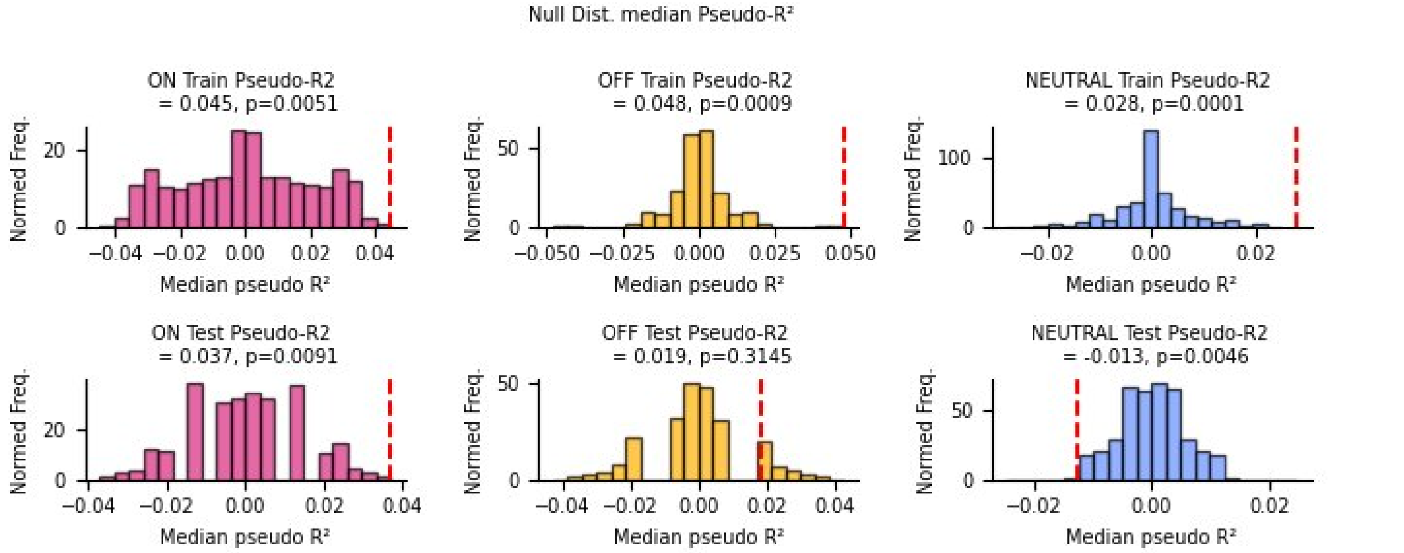


Figure S4: Two sided T-test for train and test $R^2$ (sign flipping permutation within animals, $N_{perm} = 10000$).

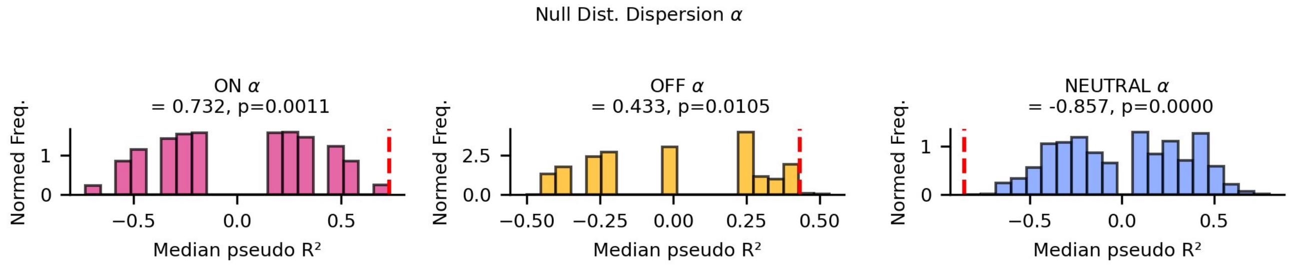


Figure S5: Two sided T-test for dispersion parameter $\alpha$ (sign flipping permutation within animals, $N_{perm} = 10000$).

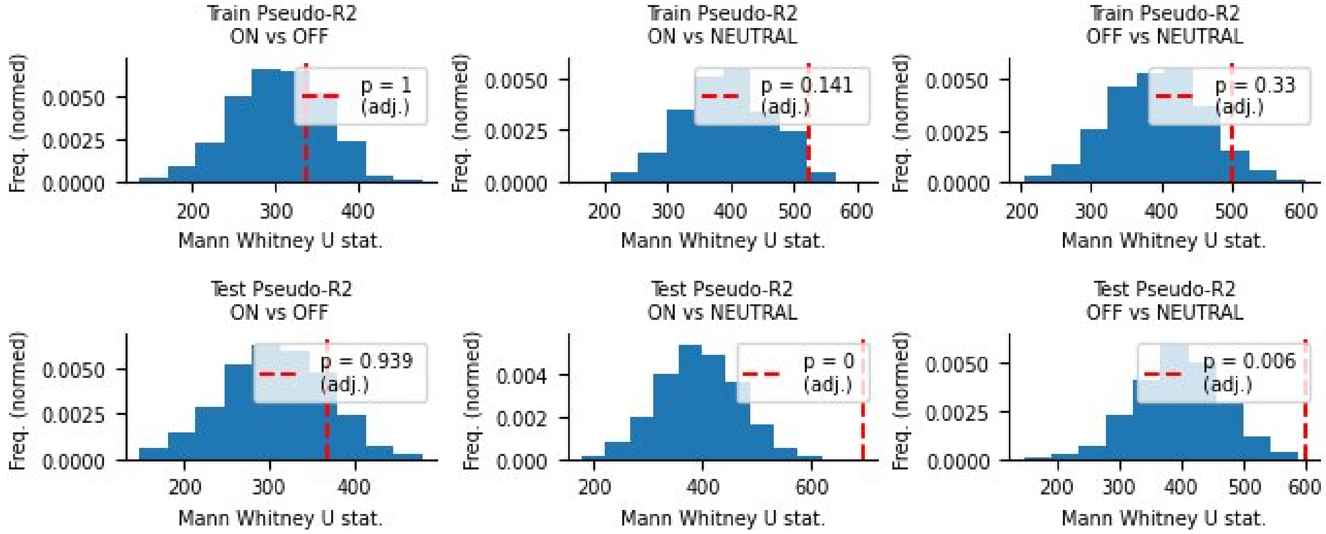


Figure S6: Mann Whitney U tests, Bonferroni corrected for multiple comparisons (sign flipping permutation within animals, $N_{perm} = 1000$).

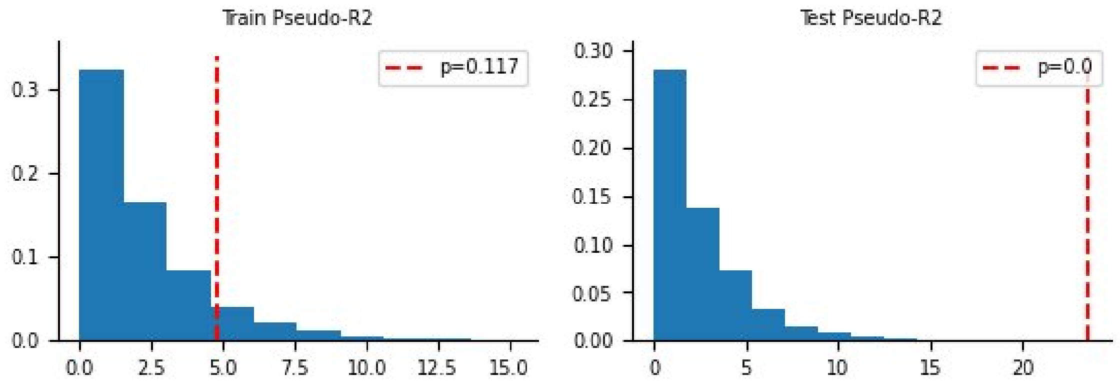


Figure S7: Kruskal-Wallis tests for differences in group medians (sign flipping permutation within animals, $N_{perm} = 1000$). Significant differences were observed at test time but not at train time.

#### Full dataset details

| Protocol | Animals | # ON-, OFF-, NEUT | Recording Details | Analyses |
|---|---|---|---|---|
| Evoked | 35 | 45, 35, 4 | 5 min. spaced trials | Piecewise Regression |
| Evoked/Ongoing | 18 | 0,0,20 | 5 min. spaced trials | Piecewise Reg., GP fit, & PSD. |
| Ongoing | 50 | 25,25,12 | Ongoing (trial bookends) | Piecewise Reg, GP fit, PSD. |
| Totals | 103 | 166 (70, 60,36) | | |

Table S11: Breakdown of data by cells recorded, trial type, and analysis performed.

#### Null Bayesian models

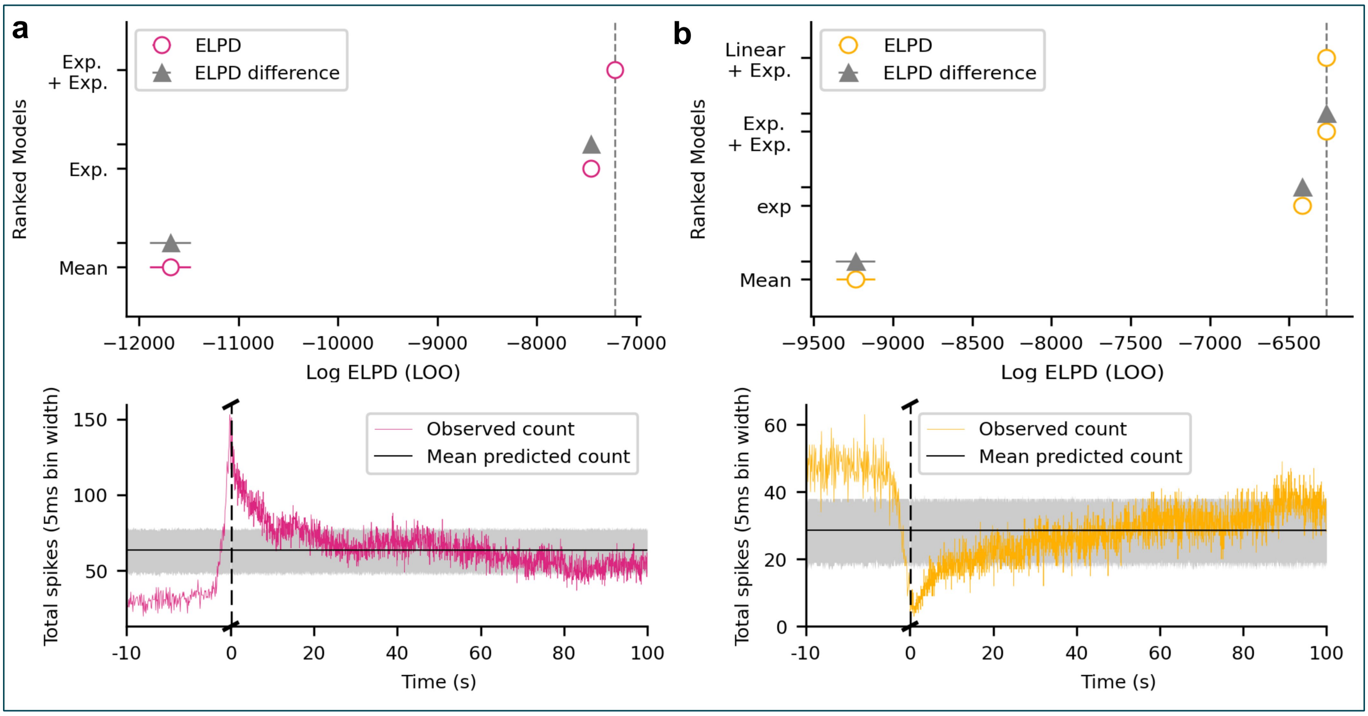


Figure S8: **a**, The null model comparison (top) and fit (bottom) for ON-cells. **b**, The null model comparison (top) and fit (bottom) for OFF-cells. Both null models clearly performed worse than either of the parametric fits.

#### Gaussian CDF Fits

The fit chosen for the OFF-cell response implies a general principle underlying the the response mechanism. We considered two different mechanisms and their associated response curves: a) A Gaussian CDF response, and b) a sigmoidal response. A Gaussian CDF response corresponds to the cumulative response of a population whose neurons cease firing in a normally distributed manner around a response time $t_0$. In contrast, a sigmoidal response satisfies the logistic differential equation:

$$\frac{df}{dx} = f(x)(1 - f(x)). \tag{9}$$

This implies that the governing population has a degree of reciprocal inhibition within it. Interestingly, we observed that the sigmoidal response function fit better to the data, despite the spiketrains being taken from independent animals over widely separated trials. The sigmoidal response function has heavier tails, the mean firing rate of the sigmoid (and hence the variance at each time given by the Poisson likelihood) remains higher for longer as the response drops towards zero than under a Gaussian CDF response (see Figure S9). This increased variance at longer response times accommodates noisier data than the Gaussian CDF, thus increasing the ELPD. Overall, we believe that the current data are insufficient to clearly distinguish between the two underlying response mechanisms, despite the increased ELPD of the sigmoidal response function.

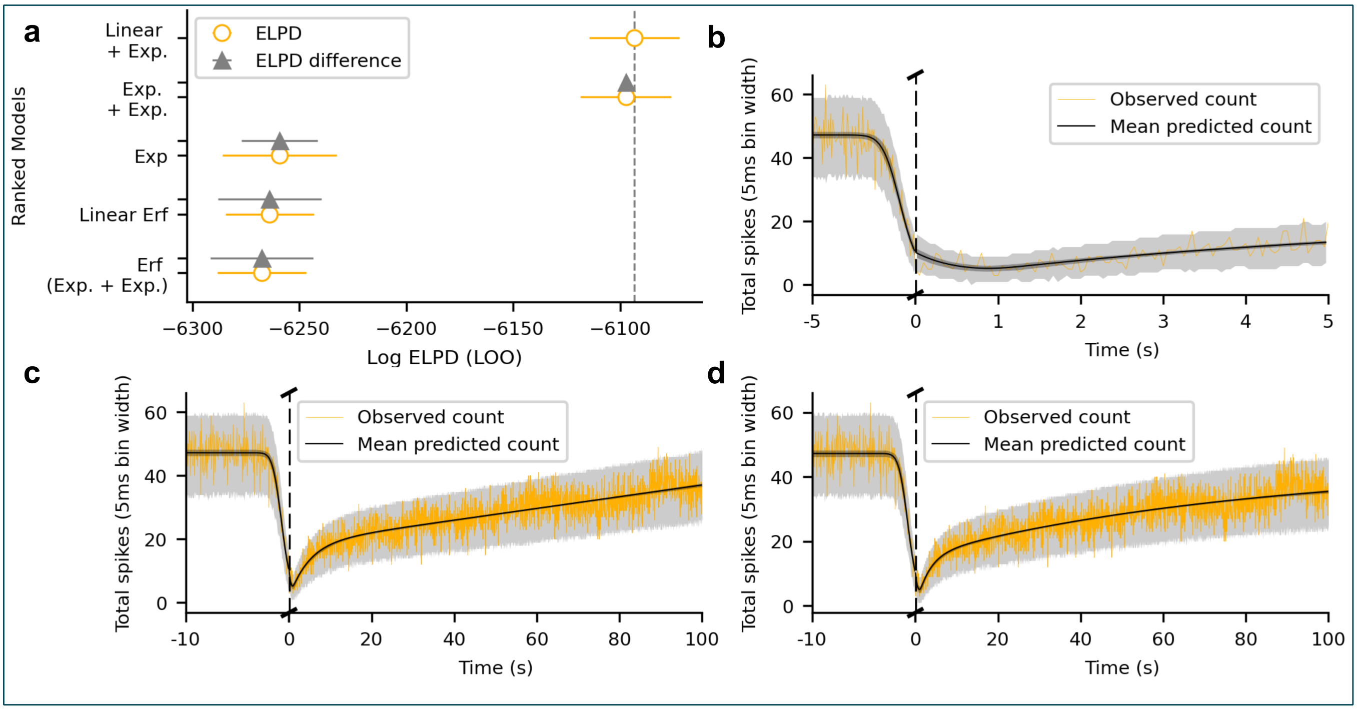


Figure S9: **a**, Model comparison of all non-baseline OFF-cell models, including the Gaussian CDF (erf) fits with a linear recovery function ("*Linear Erf*") and with a double exponential recovery ("*Erf (Exp. + Exp.)*"). **b**, A zoomed in view of the "*Linear Erf*" model fit around t=0, showing very good visual agreement with the OFF-cell decay. **c**, The full "*Linear Erf*" fit to the data, and **d**, The full "*Erf (Exp. + Exp.)*" fit to the data. Whilst the initial decay is visually identical in both, differences can be seen in the predicted recovery trajectory, paralleling those seen in the better performing sigmoidal models.

#### GP residuals

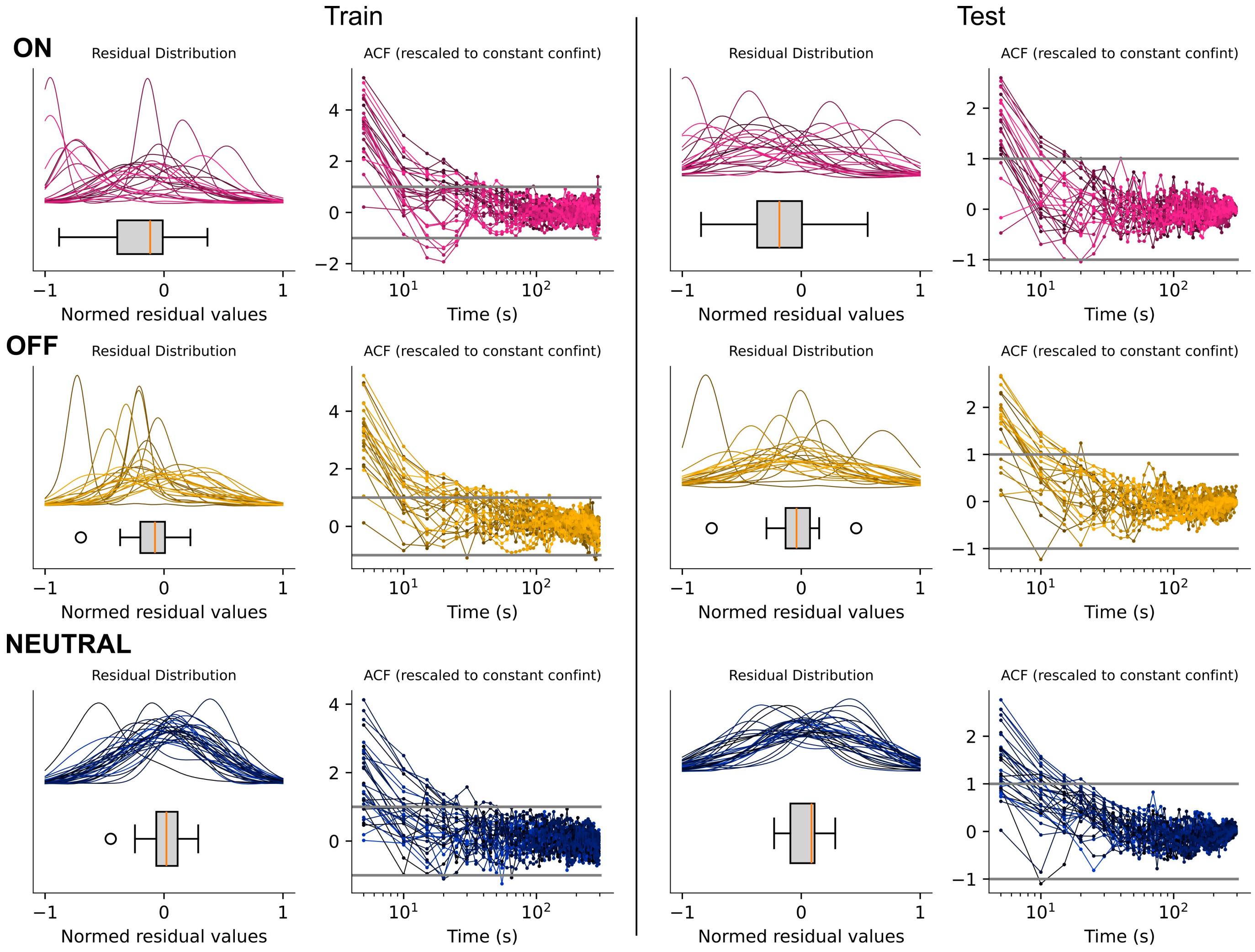


Figure S10: Gaussian process residuals for each cell. Residuals were normalized between -1 and 1 for combined plotting. Boxplots show the distribution of the mean of each residual. Fliers (circled) are Q3+1.5 IQR or Q1-1.5 IQR.

#### Truncated GP fits

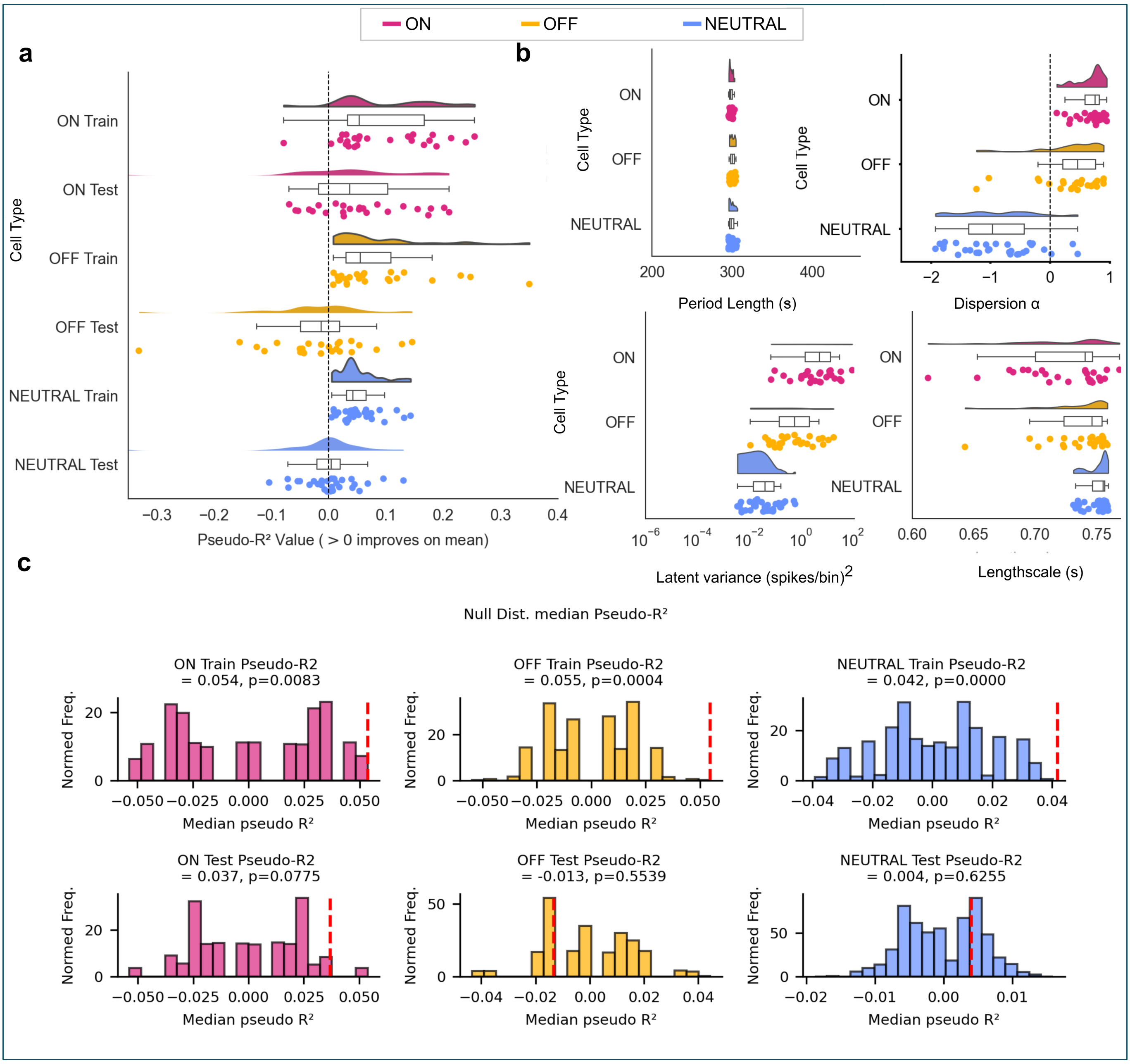


Figure S11: Results for the truncated training data. **a**, Training and test prediction accuracies. **b**, Hyperparameters of the fitted GPs. **c**, Hierarchical Pseudo-R2 tests for the truncated models.

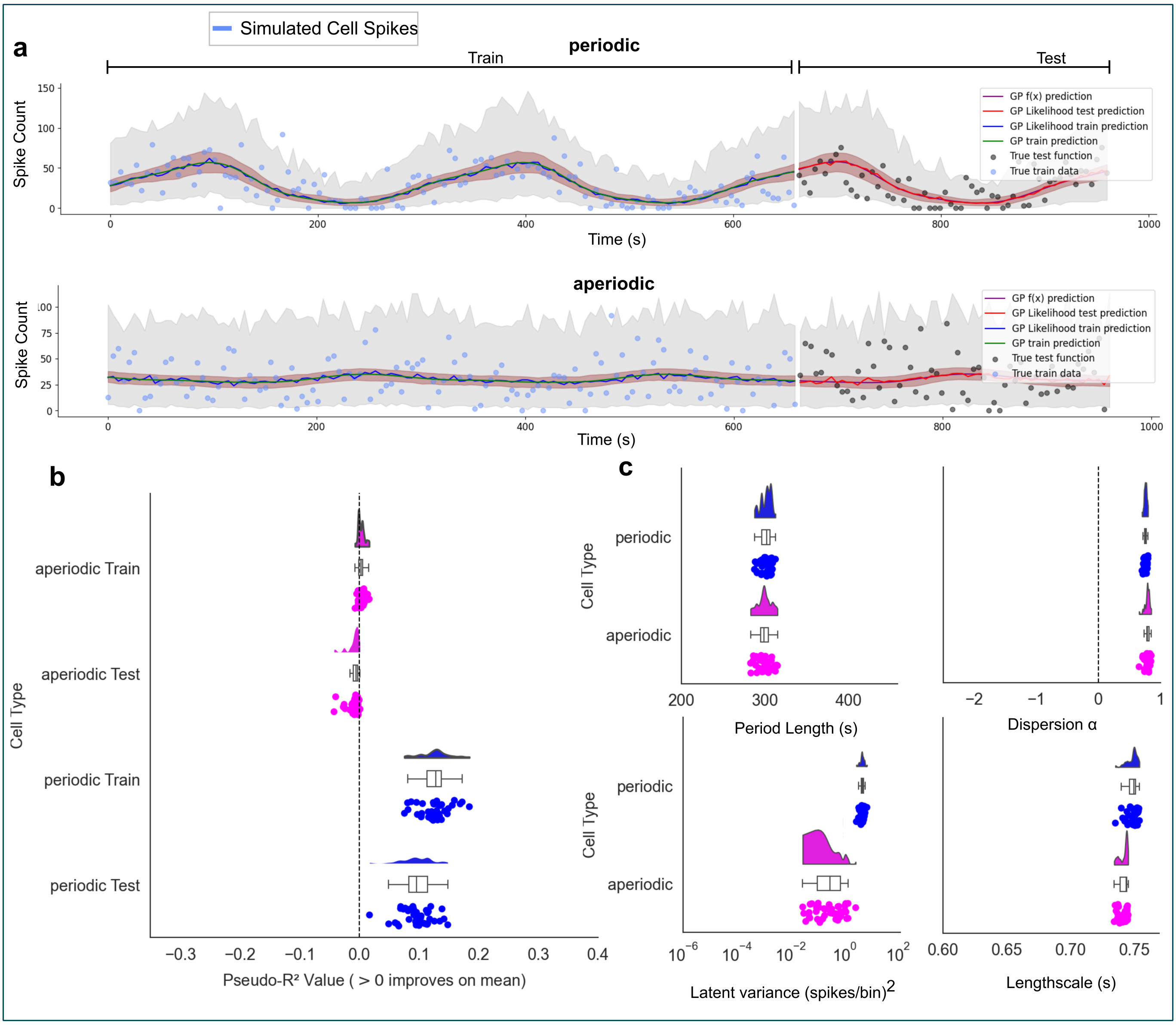


Figure S12: Results for the Poisson GP fit on a simulated periodic and aperiodic cell over 660 seconds of training data. **a**, Sample periodic (top) and aperiodic (bottom) cells. **b**, Prediction accuracies for the periodic and aperiodic cell classes, showing better than average predictive power for the periodic class and no predictive power for the aperiodic class. **c**, Hyperparameter fits for the periodic and aperiodic classes.

#### Coherence vs correlation

It is possible for two signals to have an arbitrarily low correlation whilst also having a perfect coherence, at a specific frequency. To see this, take a common sinusoid $s(t) = A\sin(\omega_0 t)$ and generate two signals $x(t) = s(t) + n_1(t)$ and $y(t) = s(t) + n_2(t)$, where $n_{1,2}(t)$ are noise processes uncorrelated with $s(t)$ which have zero power at the frequency $w_0$, for example, white noise passed through a brick wall filter (or more realistically a notch filter). Then we can compare the lag-0 cross-correlation and coherence of $x(t)$ and $y(t)$. The covariance between the signals is:

$$\begin{aligned} \mathrm{Cov}(x(t), y(t)) &= \mathbb{E}[x(t)y(t)] = \mathbb{E}[s(t)^2 + s(t)n_1(t) + s(t)n_2(t) + n_1(t)n_2(t)] && (10)\\ &= \mathbb{E}[s(t)^2], && (11)\\ & && (12) \end{aligned}$$

with correlation coefficient:

$$\rho = \frac{\text{Cov(x(t),y(t))}}{\sqrt{\mathbb{E}[s(t)^2 + \sigma_1^2]}\sqrt{\mathbb{E}[s(t)^2 + \sigma_2^2]}}. \tag{13}$$

Therefore, as $\sigma_1, \sigma_2 \to \infty$, the correlation between $x(t)$ and $y(t)$ will approach 0.

However, because both signals contain $s(t)$, the coherence at the frequency $\omega_0$ is:

$$\gamma_{xy}^2(\omega_0) = \lim_{T \to \infty} \frac{\left| S_{xy}^{(T)}(\omega_0) \right|^2}{S_{xx}^{(T)}(\omega_0)\, S_{yy}^{(T)}(\omega_0)} = 1,$$

where we take a limit as $T \to \infty$ of time limited spectra to account for the sinusoidal nature of the signal $s(t)$ having a delta function spectrum (the division of delta functions is undefined). From this we can also see that if two signals are completely correlated, then they will be coherent, but they can be coherent at a specific frequency, then they do not need to be correlated in general, if there are other processes at different frequencies influencing the two signals. Therefore, we use coherence to detect relationships between heart rate and the ON- and OFF-cell periodicity in the presence of decorrelating noise.

### Histology Images

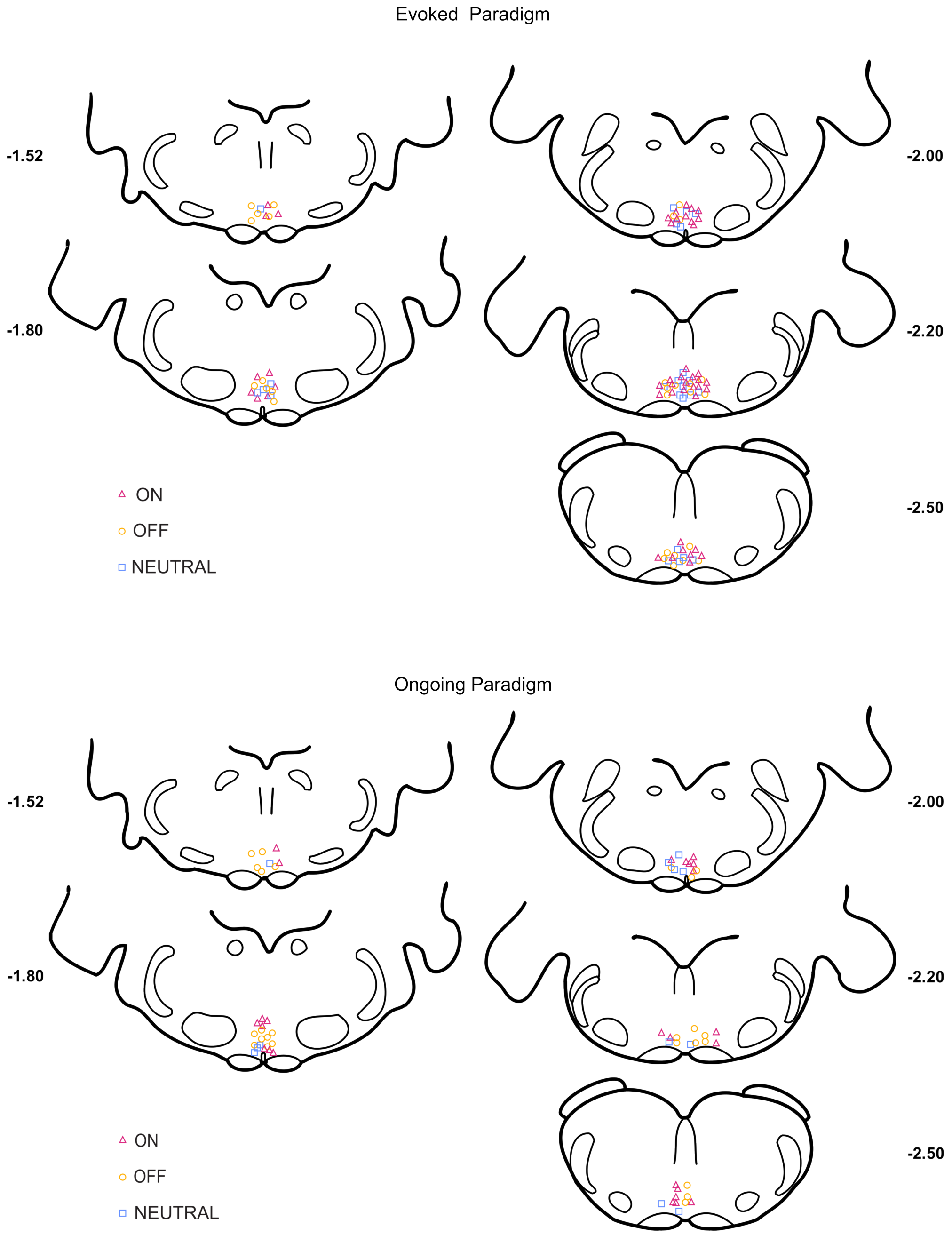


Figure S13: Reconstructed locations of all cells recorded from the RVM in the evoked and ongoing paradigms.
